# Process-based model of fish incubation survival for designing reservoir operations: a case study for Sacramento River winter-run Chinook salmon

**DOI:** 10.1101/2020.08.26.269357

**Authors:** James J. Anderson, W. Nicholas Beer, Joshua A. Israel, Sheila Greene

## Abstract

Allocating reservoir flows to meet societal and ecosystem needs under increasing demands for water and increasing climatic variability presents challenges to resource managers. Often, regulated rivers have been operated to meet flow and temperature compliance points that mimic historical patterns. Because it is difficult to assess if this approach is efficient or equitable, new more process-based approaches to regulation are being advanced. This paper describes such an approach with a model of egg incubation survival of Sacramento River winter-run Chinook salmon (SRWRC, *Oncorhynchus tshawytscha*). Thermal mortality only occurs in a critical window around egg hatching when the embryo is most sensitive to temperature stress. The duration of the critical window has significant implications for Shasta Reservoir operations that are designed to control temperature during SRWRC incubation. Previous operations sought to maintain a low temperature over the entire incubation period. However, model analysis suggests that targeting cold water directly to the critical egg hatching stage provides higher survival while requiring less cold water resources. The calibrated model is publicly accessible through a web interface connected to real-time river and fish databases and a river temperature forecast model. The system is an example of the next step of river management that integrates databases with hydrological and process-based biological models for real-time analysis and for forecasting effects of river operations on the environment.

## Introduction

A guiding principle for managing regulated rivers is allocating flow for both the needs of the ecosystem and for human society (Arthington 2015). Over several decades the focus has been on environmental flows (e-flow) for the restoration and sustainability of ecosystem services and for aquatic biodiversity. While better e-flow management is urgent to sustain the environment in a changing world, balancing competing demands is difficult (Arthington et al. 2018, Williams et al. 2019). Most progress has been made in empirically linking flows to ecosystem states such as species abundance (Chen and Olden 2017) while acknowledging the need to include greater processes or mechanistic understandings of flow-ecosystem connections (Arthington et al. 2018). Although a small percentage of studies have considered the effects of e-flow on ecological rates and, in particular, demographic processes, such as colonization and persistence of populations (Wheeler et al. 2018), in general such approaches are rare.

Exceptions to this wide ecosystem focus are found in regulated rivers in the United States that are required by the National Marine Fisheries Service (NMFS) to protect endangered species. Two prime examples are the Columbia and Sacramento rivers where management is dictated by Biological Opinions (BiOps) issued for the protection of endangered salmon (NMFS 2019a, b). These BiOps do consider many aspects of the river basins’ ecology, but the main focus is to benefit critical life-stages of salmon. For example, in the Columbia River, water spills at dams are increased to protect the juvenile salmon during their downstream migration (NMFS 2019b); and in the Sacramento River, water releases from the Shasta Reservoir are timed to cool the river during the incubation of winter-run Chinook salmon (NMFS 2019a). These salmon-centric foci have facilitated the development of process-based operations models to a degree beyond what has been achievable by the e-flow approach (Davies et al. 2014). Of note, in the Sacramento system, much work has focused on evaluating optimal reservoir operations under both drought and high water years (NMFS 2019a, Notch et al. 2020). A particular focus has been temperature, which is one of the most important factors controlling freshwater survival of salmon (Bartholow 2004, Alexander et al. 2015, Martin et al. 2017). The literature has expressed similar concerns (Olden and Naiman 2010), noting it is not sufficient in e-flow restoration to mimic natural flow regimes as we enter a future of changing natural baselines (Capon et al. 2018). The challenge was succinctly framed in a study to promote both fish and hydroelectric production in the Mekong River (Sabo et al. 2017), not as “How much water do we need?” but rather, “When do we need it most and when can we spare it?” However, answering these questions based on mimicking statistical elements of natural flows has been controversial (Williams 2018, Sabo et al. 2019). In contrast, for the Sacramento salmon the question of flow allocation has been more process-based in terms of the upper critical temperature of egg incubation.

The road to process-based management of the Sacramento River began in the 1940s with the construction of Shasta and Keswick dams on the Sacramento River. Prior to the dams, the SRWRC spawned in the cool, headwater tributaries of the Sacramento River. After dam construction, the fish spawned in the warmer water reaches between Keswick Dam and Bend Bridge (Fig. 1), so construction of a temperature control device provided improved capabilities for Shasta Reservoir operations to mimic the historical habitat with cold water releases. This remedy was judged to put the run at a low to moderate risk of extinction (Lindley et al. 2007), in particular because of the possibility of a prolonged drought depleting the cold water storage or a failure of cold water management (Williams 2006). These possibilities, predicted in 2006, were realized with the onset of California’s longest drought beginning 2011 (Seager et al. 2015) and the inability of the Shasta Reservoir operations to access the diminished cold water from the bottom of the reservoir in 2014 (Hallnan et al. 2020). As a result, for two consecutive spawning seasons (2014-2015) the incubating winter-run eggs and alevin experienced unprecedented warm waters and low survival between egg deposition and fry passage at Red Bluff Diversion Dam (RBDD) (Fig. 1). An analysis of Shasta Dam temperature operations showed that with extremely dry conditions, the thermal conditions best for Chinook salmon were unlikely to be achieved (Thompson et al. 2011, Sapin Joseph et al. 2017), although modifications to the dam could provide improvements for some species (Clancey et al. 2017).

**Figure 1.**
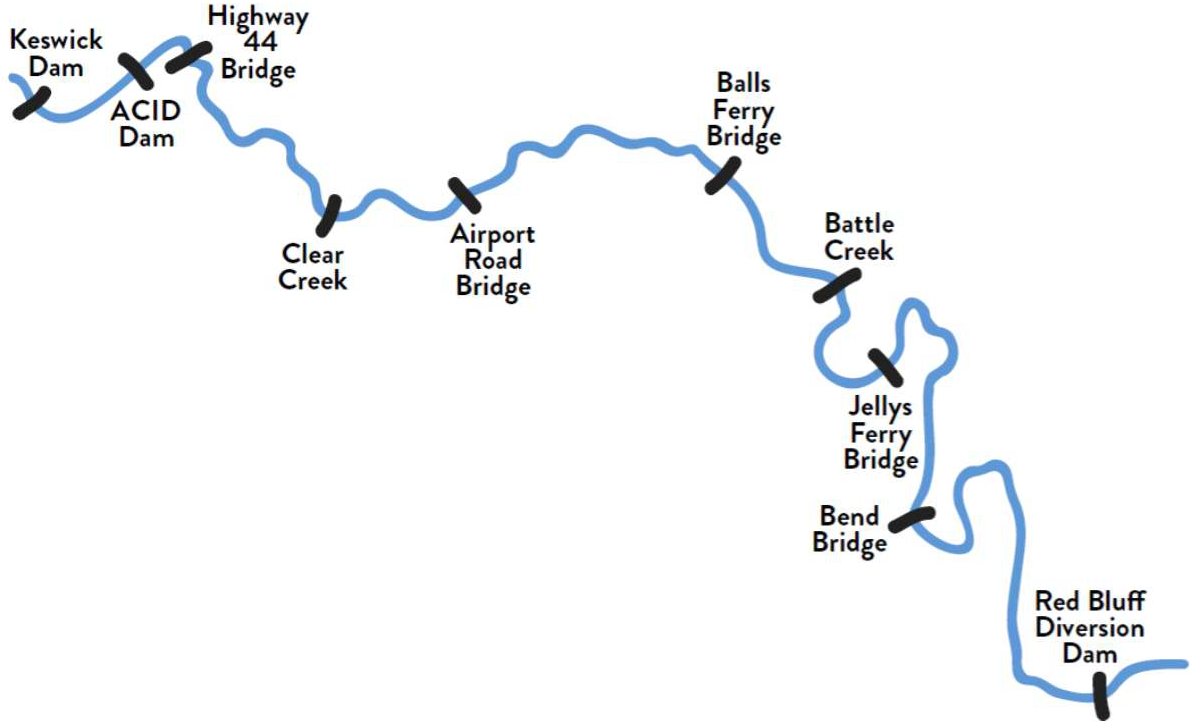
Upper Sacramento River showing reaches of winter-run Chinook salmon spawning habitat.

In 2017, the Bureau of Reclamation proposed a new long term operation plan for the Central Valley Projects and developed performance measures that link the Sacramento River temperature to the biological needs of SRWRC egg incubation. The plan specified that between May 15 and October 31 the Bureau of Reclamation (Reclamation) operate Shasta Reservoir to best utilize temperature capabilities to focus temperature management between Keswick Dam and the confluence with Clear Creek (Fig. 1). Since the ability to regulate temperature depends on the cold water volume in the reservoir, the compliance temperature would be adjusted; with 53 °F (11.7 °C) for the entire period in the best years, to a shorter window of 53 °F in some years, and in worse case scenarios meeting 56 °F (13.3 °C). Thus, if cold water were unavailable to maintain a low temperature over the period, the compliance temperature may be set higher and/or duration of these optimal temperature shortened.

The LTO plan used insights developed from a biophysical model linking mortality to river temperature (Martin et al. 2017). Importantly, the model proposed ‘stage-independent’ mortality in that temperature sensitivity occurs throughout the entire incubation period, with equivocal mortality rates during all developmental stages from egg fertilization, through hatching, emergence and passage at RBDD. However, studies indicate that the sensitivity to temperature changes as the embryo grows (Rombough 1994). An alternative model was developed (Anderson 2018) that proposed ‘stage-dependent mortality’ in which the critical temperature sensitivity occurrs during a much narrower window around the time of hatching. Prior to, and after this ‘critical window’ of development, temperature sensitivity is reduced. The final BiOp (2019a) evaluated both the stage-independent model (Martin et al. 2017) and the stage-dependent model (Anderson 2018) and proposed multiple-tier operations in which the coverage of the critical window depends on the available cold water volume.

In this paper we advance this process-based approach of the stage-independent and stage-dependent models by expressing the duration of the critical window as a parameter. We thus combine the two alternative models into a flexible form and explore the model properties, calibration, and implications to temperature management of the Sacramento River.

## Methods

### Total mortality

The model is an extension of a model developed by Martin et al. (2017) to described mortality in fish incubation in terms of the effects of temperature on respiratory demand of winter-run Chinook salmon egg incubation. The model characterizes thermal, density dependence and background mortality processes, with survivals *V, U* and *B* respectively. The total survival is *S* = *V* ⋅*U* ⋅ *B*. The model parameters described in the following sections are listed in Table 1.

**Table 1.** Definitions of model fitted parameters and symbols

| Reference | Symbol | Description |
| --- | --- | --- |
| Eq. 15 | $B$ | background survival |
| Eq. 14 | $U$ | density-dependent survival |
| Eq. 2,5,7 | $V$ | temperature-dependent survival |
| Eq. 15 | $S$ | survival from fertilization to RBDD passage |
| Eq. 13 | $L_k, k, R$ | reach length, number, total number of reaches |
| Eq. 14 | $K, D$ | carrying capacity per habitat and per km of river |
| Eq. 13,16,17 | $\rho, A$ | redds per km per reach, redds per habitat |
| Eq. 3 | $\delta$ | thermal window width in days |
| Eq. 6 | $d$ | day of year of fertilization |
| Eq. 2,4 | $y$ | embryo age in days |
| Eq. 3 | $Y_0, Y$ | age beginning and end of critical window |
| Eq. 4 | $E$ | age fry emergence |
| Eq. 6,8 | $ATU_Y$ | ATU at end of thermal window |
| Eq. 9 | $ATU_{crit}$ | ATU at maximum thermal mortality |
| Eq. 4,5 | $T_{crit}$ | critical temperature |
| Eq. 2,6 | $T_y$ | temperature at incubation age $y$ |
| Eq. 9 | $\bar{T}$ | mean temperature over incubation |
| Eq. 11 | $\Delta T_{\delta}$ | temperature differential in critical window |
| Eq. 4,10 | $b_{\delta}$ | fixed mortality rate over thermal window |
| Eq. 4 | $\beta_y$ | age-specific thermal mortality rate per day |
| Eq. 12,16 | $opt$ | differential of logit fitting |
| Eq. 10, Table 5 | $c_0$ | coefficient for $b_{\delta}$ vs $1/\delta$ regression |
| Eq. 8, Table 5 | $a_0, a_1$ | coefficients for $ATU_Y$ vs $\delta$ regression |
| Table 5 | $b_0, b_1$ | coefficients for regression of $B$ to $D$ |
| Eq. 1 | $m, m_0, m_1$ | metabolic rate, coefficients for egg metabolism |
| - | $f_k, a_k$ | population in fraction reach $k$, reach attractiveness |

### Thermal mortality

In the stage-independent model (Martin et al. 2017), mortality from thermal stress results when the embryo metabolic oxygen demand exceeds the diffusive supply of oxygen through the egg membrane. With the implicit assumption that metabolic demand increases with temperature, the mortality rate is proportional to the differential of temperature above a critical temperature. This model assumes the relationships are fixed across all stages of embryo development until fry emergence from the redd.

#### Physiological evidence of stage-dependent mortality

Rombough (1994) showed that metabolic oxygen demand increases in an exponential-like manner up to the maximum total dry mass (MTDM), then declines prior to fry emergence, as illustrated in Fig. 2 In contrast, the flux of oxygen to the organism exhibits a step-like increase upon hatching when the embryo switches from diffusion controlled cutaneous flux to branchial respiration in which oxygenated water is pumped across the gills (Wells and Pinder 1996b, a). The increased transport efficiency with hatching is reflected by the step-like drop in oxygen required to maintain normal respiration (Rombough 1986) (Fig. 3). Lower critical oxygen levels indicate higher tolerance for low oxygen levels, thus after hatching the embryo is approximately three times more tolerant to low oxygen levels than immediately prior to hatching. Additionally, the post-hatching oxygen sensitivity is largely independent of incubation temperature in Chinook salmon embryos (Rombough 1986). The greater tolerance to low oxygen after hatch is observed across a range of studies including rainbow trout (*Oncorhynchus mykiss*(Walbaum)) (Ciuhandu et al. 2005, Ciuhandu et al. 2007), steelhead (*O. mykiss*) (Rombough 1986) and Atlantic salmon (*Salmo salar*) (Wood et al. 2019).

**Figure 2.**
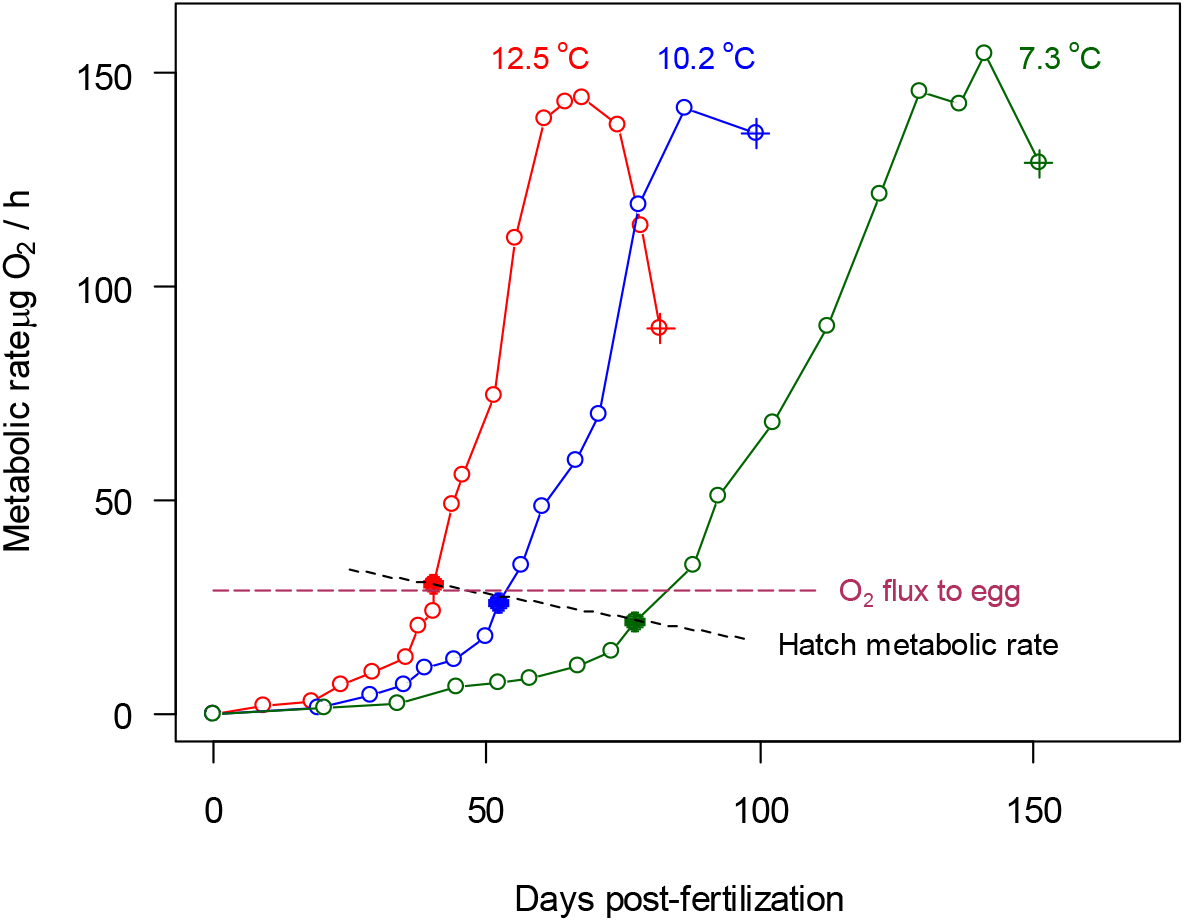
Incubating Chinook salmon metabolic rate vs. days post-fertilization at three temperatures. Redrawn from Rombough (1994). Dotted lines depict hypothetical oxygen flux to embryo in redd required to induce mortality at temperature > 12 °C. Dashed line characterizes the temperature dependence of the metabolic rate at hatch. Metabolic rate at hatching denoted (•), peak metabolic rate corresponds to the MTDM and rate at alevin emergence denoted (⊕).

**Figure 3.**
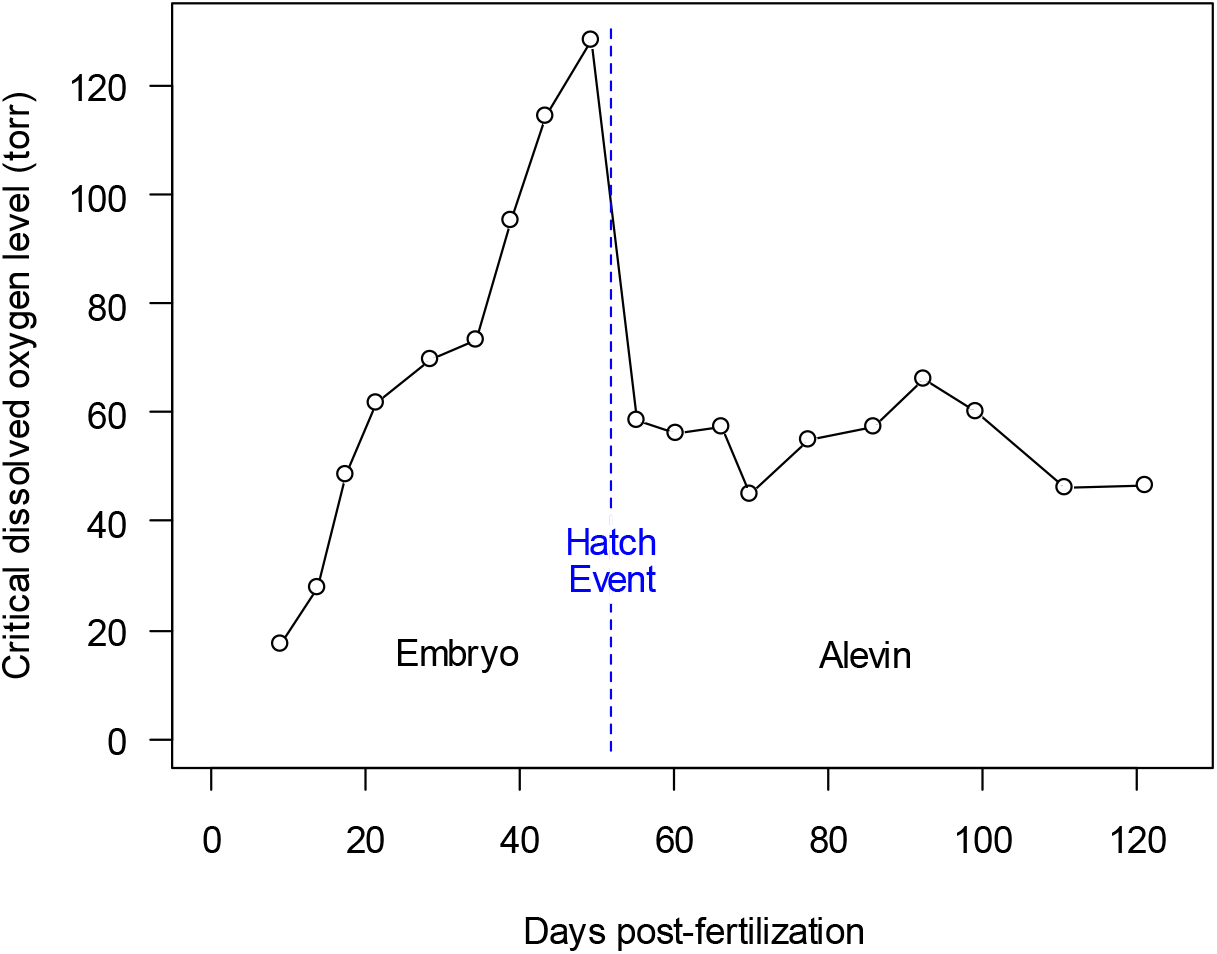
Critical oxygen as function of post-fertilization days for Chinook embryos and alevins at 10.2 °C and initial egg size 341 mg. Redrawn from Rombough (1986).

The shape of the metabolic oxygen demand with age and temperature are of particular importance for understanding the nature of stage-dependent mortality. To illustrate, consider Fig. 2 in which fish incubated at 7.3°, 10.2° and 12.5°C exhibited metabolic rates at hatch of approximately 20, 25 and 30 μgO_2_/h respectively. From (Martin et al. 2017), thermal mortality occurred above temperatures of 12°C, which suggests that the diffusive oxygen flux to eggs was between 25 and 30 μgO_2_/h, since mortality was not observed at 10.2°C.

To quantify the change in metabolism prior to hatching we determined through regression analysis that the metabolic rate, *m*, around hatching can for Fig. 2 be approximated as

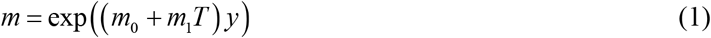

where *T* = temperature (°C), *y* = age post-fertilization (d; days) and the coefficients are *m*_0_ = − 0.0264, and *m*_1_ = 0.0087. Importantly, the metabolic rate at hatch and the daily rate of increase are both higher at warmer incubation temperatures. For example, at 12°C, with hatching on day 43, the metabolic rate is 28.6 μgO_2_/h and the rate of increase is 2.2 μgO_2_/h/d while at 10 °C, with hatching on day 53, the metabolic rate is 24.8 μgO_2_/h and the rate of increase is 1.5 μgO_2_/h/d. Martin et al. (2017) concluded that SRWRC experience incubation mortality at 12°C thus the rapid rise in metabolic rate and high rate at hatching appear critical factors inducing the onset of mortality in SRWRC. Importantly, at 12°C a change in the metabolic rate from a safe level of 24.8 μgO_2_/h to a mortality-inducing level of 28.6 μgO_2_/h can occur over a few days.

A number of studies suggest the effect of low oxygen on post-hatch growth, respiration and survival is less pronounced (Rombough 1986, Ciuhandu et al. 2005, Ciuhandu et al. 2007, Wood et al. 2019). Figure 2 illustrates that respiration was unaffected by temperature at MTDM for Chinook salmon and Fig. 3 illustrates that the oxygen sensitivity is unchanged after hatching. Additionally, laboratory studies with SRWRC salmon indicate that the critical temperature for alevin is above 13.3 °C (USFWS 1999). Thus, fish surviving the oxygen levels encountered at hatching are likely to survive those same levels in the alevin stage. Little information is available on effects of oxygen on alevin movement but studies indicate that coho (*Oncorhynchus kisutch*) larvae moved vertically tens of centimeters after hatching and the initial depth of burial and gravel size did not affect alevin survival or timing of emergence (Dill and Northcote 1970). Taken together, these studies suggest that a few days preceding hatching is the most critical period for thermal mortality. We designate this period the critical window.

#### Thermal mortality rate

To model mortality over the critical window first express survival from thermal stress as

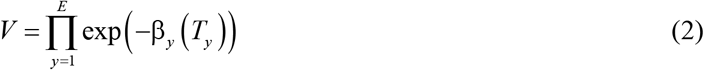

where β_*y*_ is the daily mortality rate at age *y* (days since fertilization) as a function of temperature *T*_*y*_ and *E* is the age of fry emergence from the redd. Note, that as is observed in laboratory studies (Rombough 1986, Rombough 1994), the mortality rate changes daily even at with constant temperature. Thus, when temperature also changes over time the change of β_*y*_ is complex. However, we can suitably approximate the pattern by assuming that the embryo thermal stress only occurs within a critical window of development between ages *Y*_0_ and *Y*. The duration of the critical window in days is

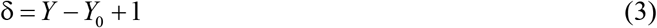

where the +1 accounts for the fact that if *Y* = *Y*_0_ the window duration is one day. Inside the critical window we characterize the mortality rate in terms of the differential in temperature above a critical threshold similar to the function used in the stage-independent model (Martin et al. 2017). The sum of daily mortalities within the window for one redd is then

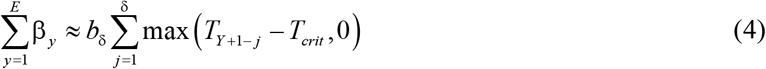

where *T*_*crit*_ is the critical temperature for thermal mortality and *b*_δ_ is a fixed mortality rate coefficient, which depends on δ. Stage-dependent thermal mortality then depends on window duration, δ, the daily temperatures *T*_*j*_ for *j* = *Y*_0_ to *Y*, and the mortality rate coefficient giving

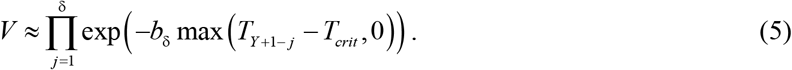

To characterize the end date of the critical window we note that that the stages of development, e.g. cleavage embryo, hatch and fry emergence (Alderdice and Velsen 1978, Beacham and Murray 1990, USFWS 1999) closely track the accumulated temperature units experienced by the embryo since egg fertilization. Furthermore, physiological studies discussed above indicate that the stress from low oxygen and high temperature increases as a unimodal peak near the hatch, and so we reason that the critical window contains this peak and characterize the critical stage of development in terms of a critical accumulation of temperature units (ATU) denoted *ATU*_crit_. Because ATU is linked to physiology the *ATU*_crit_ is the same for all redds but the day-of-year at which it occurs depends on date of fertilization and the unique temperature experience of each redd. The end date of the critical window is determined by the ATU experienced since fertilization on day-of-year *d* as

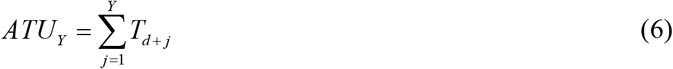

If *Y* = *E* and *Y*_0_ = 1, the critical window is the entire incubation period, i.e. δ = *E*, and Eq. (5) is identical to the thermal mortality in the stage-independent model (Martin et al. 2017). For a minimum critical window of one day, i.e. δ = 1 d, the thermal survival equation reduces to

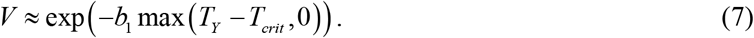

where the day-of-year of thermal mortality is then *Y*_0_ = *Y*.

#### Relationship of model parameters with *δ*

The critical window width δ is a central parameter in the model that *a prior* can fall anywhere between one day and the entire incubation period of a redd. Because of its importance in determining how much and when to target cold water during incubation we consider δ from statistical and biological perspectives. The sections below give results and Appendix S1 gives the derivations of Eqs. (8) to (12).

#### *Relationship of ATU to* δ

Ultimately the critical window is defined by the stage of development of the embryo as characterized by the ATU over incubation. The ATU at the end of the window, *ATU*_Y_, increases linearly with the window width as

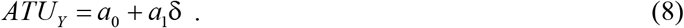

and middle of the window, *ATU*_crit_, is

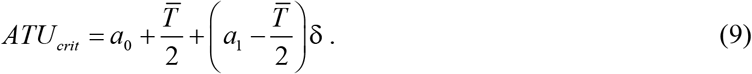

where 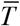 is the mean temperature the embryo experiences since fertilization.

#### *Relationship of b*_δ_ *to* δ

The intrinsic rate of thermal mortality completely depends on the selection of critical window width because to fit an observed mortality the daily intrinsic rate must be large when all the mortality is confined to small window and the daily rate must be small when mortality is spread over a large window. The relationship is

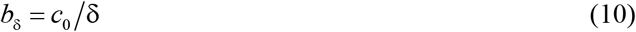

with *c*_0_ = *b*_1_ Δ*T*_1_/Δ*T*_δ_ where, over the window, the temperature differential characterizing the mean thermal stress is

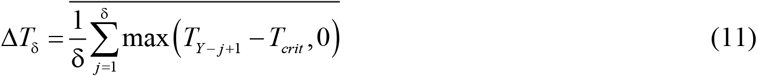

If Δ*T*_δ_ is independent of δ then *c*_0_ = *b*_1_. As is developed in the results section, this equivalence results because *b*_1_ determined by fitting the model with δ = 1 equals c_0_ determined from a zero-intercept linear regression of Eq. (10) for *b*_δ_ fit for δ 1 to 80 d. As an aside, management operations of Shasta Reservoir have sought to maintain a stable river temperature during fish incubation. Therefore, a near constant Δ*T*_δ_ has been facilitated by reservoir operations.

#### *Relationship of opt to* δ

The relationship of the model fitting error to the critical window width gives further information on plausible window widths. The relationship between the fitting error, *opt*, and the differential between a trial window width δ and the best fitting window width δ* is

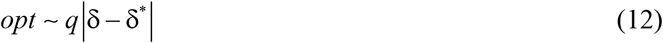

where *q* is proportional to the slope of the relationship of δ to *opt*. Over the range of possible window widths, δ = 1 to 80, the slope can change reflecting how well the thermal stress index Δ*T*_δ_ derived for δ characterize the best fitting thermal stress Δ*T*_δ_* derived from δ*. For ranges of δ in which *q* ≈ 0 the variations of temperature within the window do not significantly add noise to Δ*T*_δ_ For other ranges of δ, the series of temperatures contribute noise to the characterization of the stress differential and therefore *q* > 0. In other words, when a selected window width includes temperature information not reflecting the thermal stress the fitting error increases.

### Density-dependent mortality

As in the stage-independent model (Martin et al. 2017), density-dependent mortality is expressed by a Beverton-Holt function (Beverton and Holt 2012) giving survival from density-dependent processes as *U* = 1 /(1 + *A/ K*) where *A* is the spawning population and *K* is the carrying capacity of the habitat at which population-dependent survival is one-half. This function describes the incremental effects of increasing numbers of redds on decreasing juvenile survival. Importantly, while *A*, and therefore *K*, are typically habitat level measures, in river environments spawning numbers can also be expressed in terms of density such as redds per unit river length. In turn, expressing population density per length allows a more accurate representation of density-dependent mortality. Additionally, correlations of redd density with stream physical properties are useful because the density can be correlated with environmental properties that contribute to habitat quality (Brumbaugh and Coleman 2017).

For our model the density of redds experienced by an individual redd *i* is defined

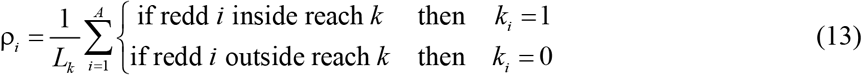

where *A* is the number of redds in the total habitat, *k*_*i*_ is the reach location index of redd *i* for reaches *k* = 1…. *R* where *R* is the total number of reaches covering the habitat and *L*_*k*_ is length of reach *k* (Table 2). In essence, this accounting divides the total habitat into *R* segments and the number of redds in each reach is counted. The density-dependent survival due to carrying capacity for each redd is then

**Table 2.** River reach indices, reference locations, mid-reach location, and reach length.

| $k$ | River Reach | Location (km) | $L_i$ (km) |
| --- | --- | --- | --- |
| 1 | Keswick Dam to A.C.I.D. Dam | 483 | 5.5 |
| 2 | A.C.I.D. Dam to Highway 44 Bridge | 478 | 3.3 |
| 3 | Highway 44 Bridge to below Clear Creek confluence | 474 | 7.7 |
| 4 | Below Clear Creek to Balls Ferry Bridge | 466 | 24.8 |
| 5 | Balls Ferry Bridge to Bend Bridge | 441 | 30 |
| Total | Keswick Dam to Red Bluff Diversion Dam |  | 94 |

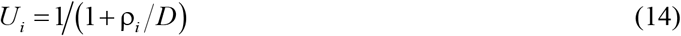

where *D* is the carrying capacity expressed in redds per km. The carrying capacity also can be expressed as a habitat level by setting *R* = 1 such that the habitat has one reach of length, *L*. Reach and habitat level measures are related as 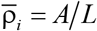 and *D* = *K* / *L* giving 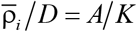.

### Background mortality

Any additional mortality that is not associated with temperature or density-dependence is designated background mortality. Although no specific mechanism is ascribed to the process, it plausibly includes mortality on fry prior to their arrival at RBDD. Background survival, which is a constant in the model, is designated *B*.

### Total survival

Total survival is expressed as the weighted sum of survivals from all redds (*i* = 1… *A*) as

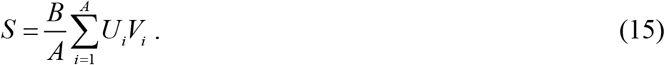

### Model configurations

A central goal of this paper was to consider different critical window hypotheses in the context of observations of incubation survival and laboratory studies of metabolism over the stages of incubation. To this end, we examples that characterize the range of possible critical window widths. Additionally, we included cases that mimic the Martin et al (2017) stage-independent egg incubation model. The models differ by the window width and spatial/temporal aggregations of redd formation (Table 3). Case 0 and Case 1 have the same temporal/spatial aggregations and a comparison demonstrates that our model structure can mimic the stage-independent model (Martin et al. 2017). Cases 2, 3 and 4, aggregate redds at the same temporal/spatial scales. Case 2, represents stage-independent thermal mortality and Case 3 stage-dependent thermal mortality. Case 4 characterizes window width between the boundaries set by Cases 2 and 3. Case 5, is stage-dependent with explicit δ = 1 d and a mean annual spawning dates, characterizes the properties of mortality by year and reach.

**Table 3.**
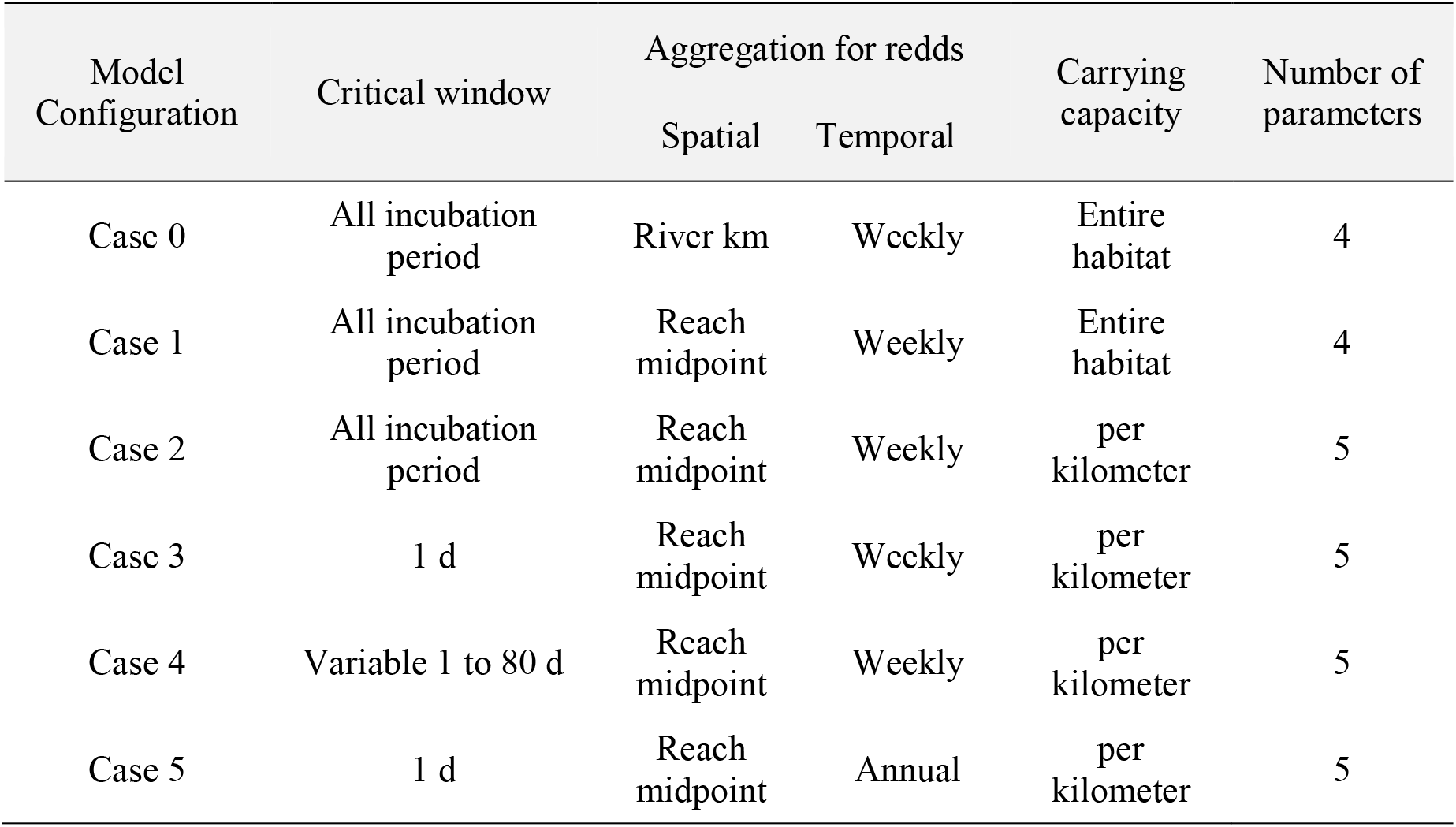
Aggregation of data and process for the model results in three configurations.

### Data

Two datasets of SRWRC spawning location/date and survival to RBDD were used in the analysis (Fig. 1). Dataset A, consisting of years 2002-2015, was used for the primary calibration and analysis. The range of data is similar to the range used in the stage-independent model. Dataset B, which includes the additional years 1997-1999 and 2017-2018, was used to explore the pairwise relationships of model parameters for differing values of δ. Survival data were not available for years 2000 and 2001 and survival from 2016, an outlier year with extremely low numbers of redds, was not used. However, including this data point does not significantly affect parameter estimates (Appendix S2: Table S2).

Temperatures were obtained from the data site www.cbr.washington.edu/sacramento/data/, which were in-turn sourced from the California Data Exchange Center http://cdec.water.ca.gov/. Daily, location-specific temperatures were calculated by linear interpolation between Keswick Dam (KWK) and Clear Creek (CCR) temperature monitors (Fig. 1). Redd count data were obtained from California Department of Fish and Wildlife aerial redd surveys (2017 Aerial redd Sacramento River summary as of 9-27-2017, from Doug Killam CDFW). Survival from egg fertilization to fry passage at RBDD was obtained from USFWS (Poytress 2016, Rea 2019).

### Model fitting

Model parameters were estimated with fits to logit-transformed-survival data of SRWRC juvenile salmon passing RBDD as a function of observed redd creation dates and incubation temperatures. Parameters were estimated with packages in the R^©^ programming language (R Core Team 2018). The Cases differ by the spatial/temporal aggregation for redd formation and the case-specific criteria by which a critical window was selected (Table 3).

Case 5, with an annual spawning scale and implicit one-day critical window, was the written in the R language and parameters were estimated with general-purpose optimization function *optim*. Cases 1-4 survivals were calculated with the SacPAS Fish Model (www.cbr.washington.edu/sacramento/fishmodel/) (Appendix S3, Fig. S1). In this form the parameter space is search and fitting criteria calculated for trial parameters generated by the derivative-free Nelder-Mead simplex algorithm *nmkb* in the *dfoptim* package. The *nmkb* routine requires upper and lower boundaries for the parameter search and used a fixed value of δ for each trial fit. Both fitting algorithms minimized the logit differential

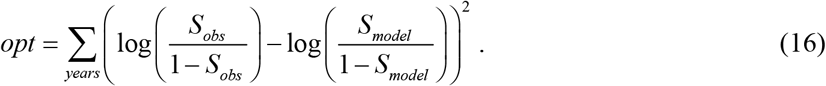

## Results

### Model fits

Results from Cases 0, 1, 2, 3 and 5 are given in Table 4. Case 0 parameters were obtained from Martin et al. (2017), which was fit with data from 1996-2015. The model has a habitat-wide carrying capacity and a critical window equivalent to the entire incubation period. Case 0 carrying capacity, *K*, was reported in spawning carcass counts and for comparison to our results was converted to redds using the ratio carcass/redd = 8.56 estimated from the geometric mean of the ratio of carcasses to redds over the years 1996-2015 (Poytress 2016). For Cases 1-3 the parameters were similar for datasets A and B. Case 1 is functionally equivalent to Case 0 with small differences due to the different years in the two datasets. Case 1 critical temperature and thermal mortality rate were slightly higher than for Case 0. The carrying capacity *K* was 3.3% less, but in compensation the backbround survival *B* was 9.0% greater. Case 2, with δ = 79 d, represents stage-independent thermal mortality and Case 3, with δ = 1 d, represents stage-dependent thermal mortality. Case 4 was used to generate the parameter pair plots for values of δ between 1 and 80 d (Fig. 4). Cases 0 and 1 have lower background survivals, *B*, than obtained for Cases 2-4. The intrinsic mortality rates, *b*_δ_, is significantly different for Cases 1and 2 compared to 3 and 4. Similarly *ATU*_Y_ is different between 2 vs 3 and 4. The basis of these differences are explained below.

**Table 4.** Parameters of model cases in Table 2 for fits to dataset A and B. R-sqr. from linear regressions of model survival against data. Case 0 parameters from (Martin et al. 2017). Cases 0, 1 and 2 represent stage-independent thermal mortality with critical window extending over entire incubation period. Cases 3 and 5 represent stage-dependent thermal mortality.

| Case | Data<sup>1</sup> | $\delta$<br>days | $B$<br>- | $T_{crit}$<br>°C | $b_{\delta}$<br>1/°C d | $D$<br>redd/km | $K$<br>redd | $ATU_Y$<br>°C d | r-sqr | $\Delta AIC^2$ | $\Delta AIC^3$ |
| --- | --- | --- | --- | --- | --- | --- | --- | --- | --- | --- | --- |
| 0 | - | 79<sup>4</sup> | 0.366 | 12.0 | 0.024 | - | 1064<sup>5</sup> | - | 0.768 | 9.3 | - |
| 1 | A | 79<sup>4</sup> | 0.399 | 12.14 | 0.026 | - | 1028 | - | 0.746 | 8.9 | 2.9 |
| 1 | B | 79<sup>4</sup> | 0.387 | 12.15 | 0.026 | - | 1137 | - | 0.724 | 9.5 | 2.7 |
| 2 | A | 79 | 0.498 | 12.15 | 0.0271 | 43.21 | - | 944 | 0.837 | 4.8 | 4.9 |
| 2 | B | 79 | 0.476 | 12.21 | 0.0291 | 44.33 | - | 929 | 0.863 | 4.7 | 2.4 |
| 3 | A | 1 | 0.487 | 12.00 | 2.24 | 43.95 | - | 488 | 0.883 | 0 | 1.4 |
| 3 | B | 1 | 0.473 | 12.04 | 2.27 | 43.11 | - | 482 | 0.895 | 0.9 | 0 |
| 5 | A | 1 | 0.514 | 11.87 | 2.20 | 38.33 | - | 550 | 0.923 | - | - |

Note 1: data extracted from (Martin et al. 2017)
Note 2: $\Delta AIC$ based on data 2002-2015
Note 3: $\Delta AIC$ based on data 1997-2018
Note 4: Equivalent $\delta$ for full incubation period critical window
Note 5: Conversion based on carcasses/redd = 8.56

**Figure 4.**
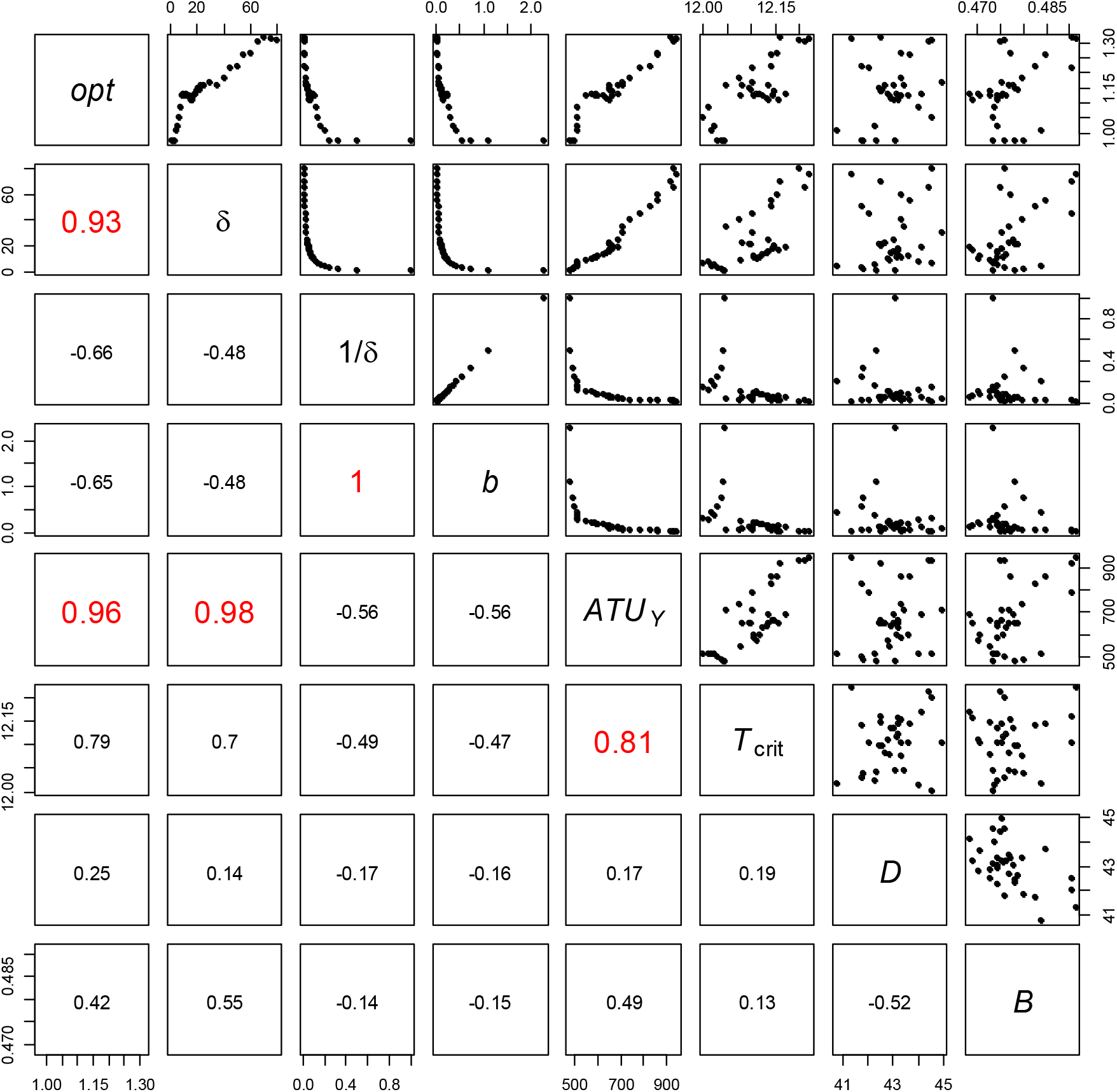
Pairwise comparisons of fitted model parameters for trial values of δ from 1 to 80 d. Upper panels show pairwise plots of model parameters for Case 4 using dataset B. Lower panels give correlation coefficients. Terms are defined in Table 1. Bolded red numbers designate highly correlated parameter pairs.

The ΔAIC scores and r-squares are calculated for datasets A (Table 4). Case 5, which ascribes all spawning to the years-specific mean spawning date with redd location set to the middle of each reach, yields the best fit. However, collapsing redd information to single locations and dates reduces the real variability in the system. We used this case to explore the interannual and reach-specific mortality processes. Effects of stage-independent vs stage-dependent thermal mortality assumptions on Shasta Reservoir temperature control operations were evaluated with Cases 1 (stage-independent mortality) and Case 3 (tage-dependent mortality).

Figure 4 presents pairwise plots of Case 3 fitted parameters obtained for critical window widths between 1 and 80 d based on dataset B. Tabular results are given in Appendix S2:Table S1. Appendix S2 Fig. S1 shows the pairwise comparison of the model fitted to dataset A. From these plots and the model theory developed in Appendix S2 we derived relationships of the model parameters (Table 5).

**Table 5.** Linear regressions of parameters derived from pairwise comparisons of fits to datasets A and B. First two regressions are based on Eqs. (10) and (8). Third regression is empirical. Setting δ = 1 in Eq. (8) *ATU*_Y_ is by definition equivalent to *ATU*_crit_.

| Regression | Dataset | intercept | slope | r-square | $c_0$ | $ATU_{crit}$ |
| --- | --- | --- | --- | --- | --- | --- |
| $b_\delta = c_0/\delta$ | A | - | 2.242 | 0.998 | 2.242 | - |
|  | B | - | 2.247 | 0.999 | 2.247 | - |
| $ATU_Y = a_0 + a_1 \delta$ | A | 492 | 5.89 | 0.937 | - | 498 |
|  | B | 512 | 5.99 | 0.951 | - | 518 |
| $D = a_2 + a_3 B$ | A | 0.649 | -0.0037 | 0.843 | | |
|  | B | 0.611 | -0.0031 | 0.275 |  |  |

First, the critical window width δ was inversely related to the thermal mortality rate *b*_δ_ as predicted by Eq. (10). Then, from a regression of *b*_δ_ vs 1/δ we obtained *c*_0_, which was found to be identical for the two data sets Furthermore, for both A and B datasets the ratio *c*_0_ *b*_1_ = 1, from which we conclude that Δ*T*_δ_ was independent of δ. This result is important because it indicates that *b*_1_ is a fundamental parameter and the rate parameter for any assumption of window width can be calculated precisely. This reconciles the estimate mortality rate determined by (Martin et al. 2017), designated *b*_*T*_ in that study, (see *b*_δ_ for Case 0 vs Case 1 in Table 4). The relationship also is the basis for exploring effects of δ on the *ATU*_crit_ and *opt*.

Second, *ATU*_Y,_ characterizing the upper boundary of the window, was linearly related to δ as predicted by Eq.(8). Using the regression coefficients from Table 5 and Eq. (9) for datasets A and B the *ATU*_crit_ is 498 and 519 °C d respectively. Additionally, with δ = 1 *ATU*_crit_ = *ATU*_Y_ then from Table 4 the *ATU*_crit_ for Case 3 using A and B datasets is 488 and 482 °C d. The average of these four estimates gives *ATU*_crit_ = 496 ± 14 °C d. With an average temperature of 11 °C the standard deviation in *ATU*_crit_ is to less than 2 days. In comparison, the ATU for Chinook hatching under laboratory temperatures similar to that of SRWRC incubation (Alderdice and Velsen 1978) is 551 ± 122 °C d.

Third, the plots of *opt* with δ exhibit a number of linear intervals that provide information of the properties of the critical window (Fig. 4, Appendix S2: Fig. S1, Fig. S2). For both datasets the *opt* is constant for δ < 5 d, increases linearly to δ ∼ 10 d, flattens between 10 and 18 d and increases gradually thereafter. For both datasets the minimum *opt* occurs for δ ≤ 5d and from Eq. (12) the flat slope indicates *q* ≈ 0, suggesting that temperature in this range of critical windows similarly characterize embryo stress. For critical windows with δ > 5 the *q* > 0 and the effect of temperature for these larger window contributes noise to the temperature differential. Presumably, the increasing *opt* at larger δ involves including temperatures in Δ*T*_δ_ corresponding to embryo ages insensitive to temperature stress.

Parameters *D* and *B* are negatively correlated (Fig. 4, Appendix S2: Fig. S1) (Table 5), which minimizes the effect of their variations on the non-thermal mortality estimate. Using this correlation the non-thermal survival can be expressed *UB* = (*a*_2_ − *a*_3_*D*) (1 + ρ *D*) where ρ is redd density. Figure 5 illustrates that survival varies little with variations in the carrying capacity parameter, *D*.

**Figure. 5.**
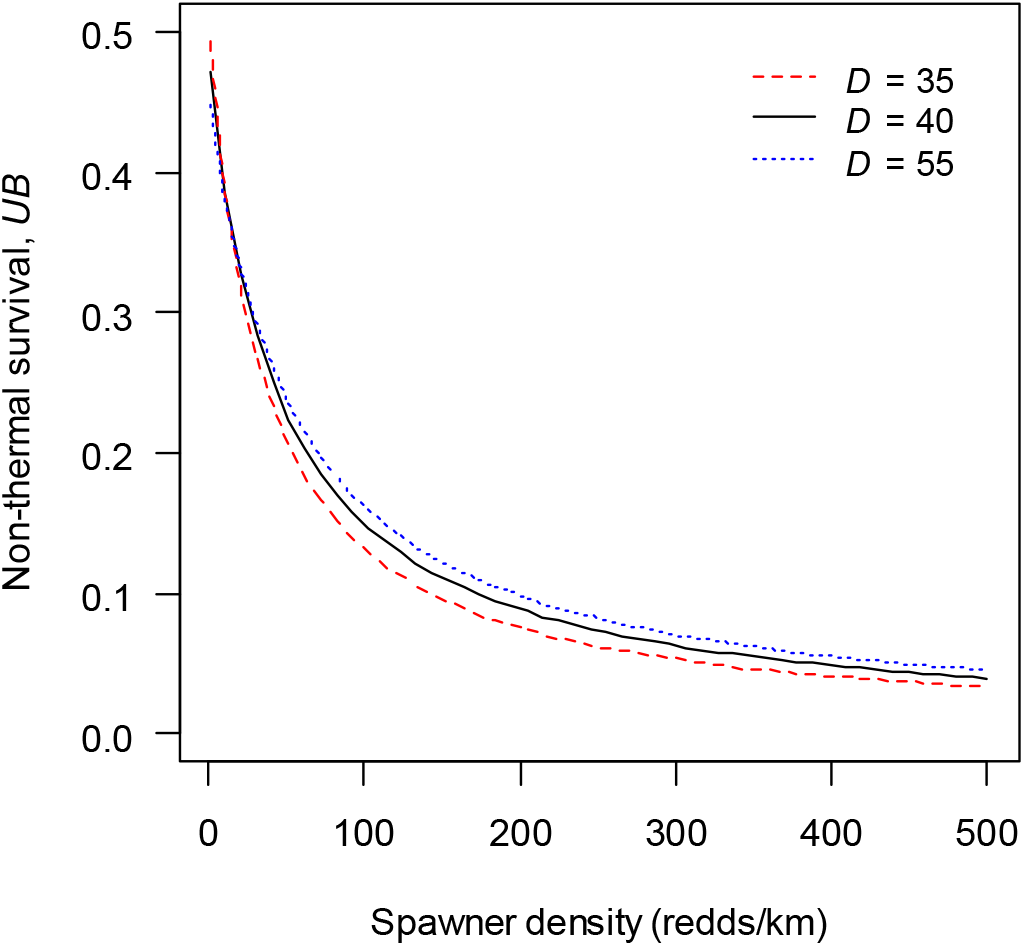
Survival from background *B* and density-dependent *U* processes for different spawner densities at three carrying capacities *D* (redds/km). Curve based on the relationship of *B* and *D* in Table 5.

### Survival patterns across years and reaches

Figure 6 illustrates the model-predicted survival to RBDD for Cases 0, 1 and 3 (Table 4). Case 3 captures range of observed survivals with 1-day thermal window. The fit is equivalent for δ < 5. The poorest fits, exemplified by Cases 0 and 1, result with the assumption of stage-independent mortality, i.e. δ = 79 d. The open points depict the survival form dataset A. The solid points depict the additional survivals contained in dataset B. Notice that all of the cases poorly predict survival for 2016. However, including this data point does not measurably change the estimated model parameters but degrades r-square (Appendix S2: Table S2).

**Figure 6.**
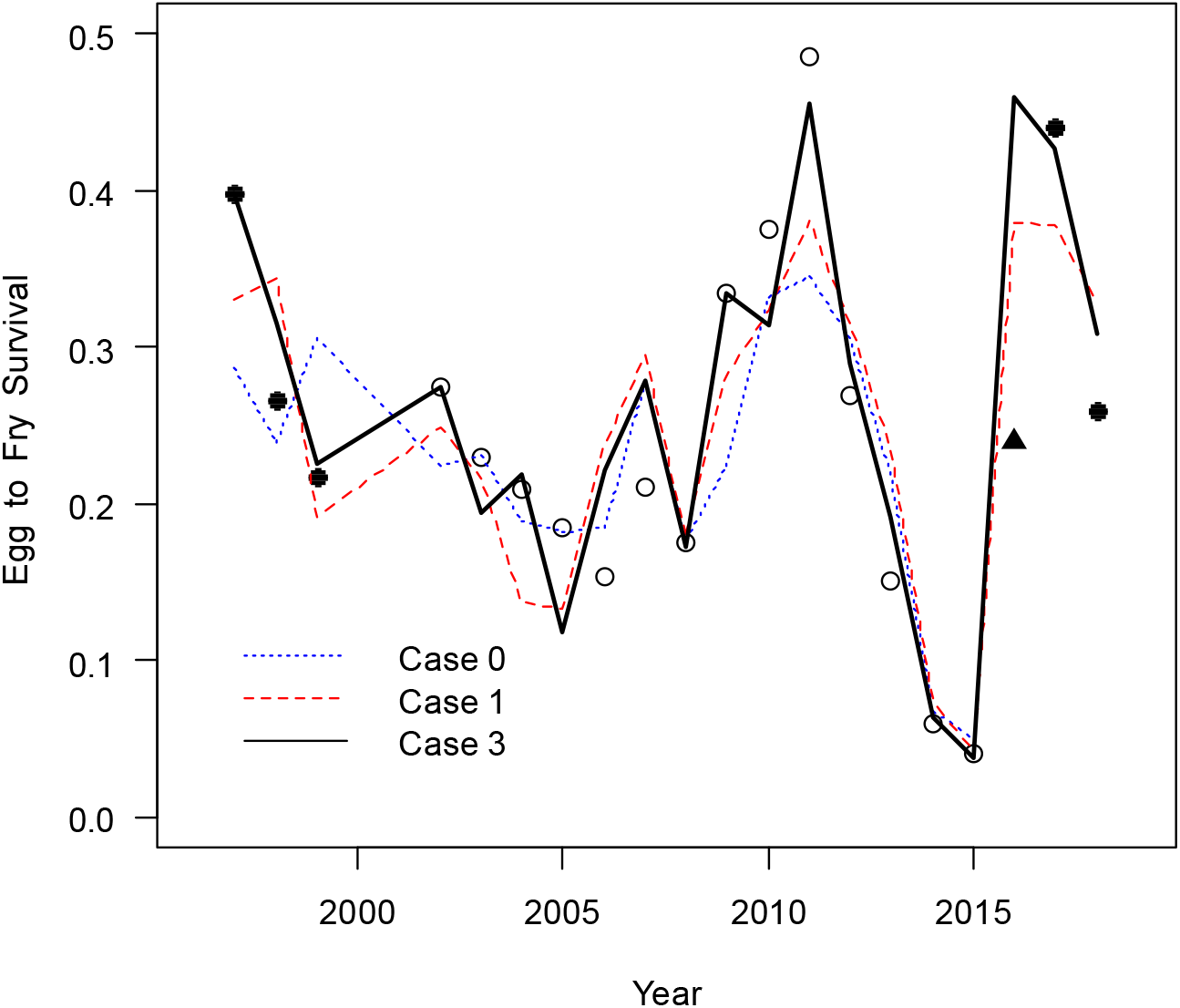
Comparison of survivals for three Cases: 0 (NMFS model), 1 (stage-inependent thermal mortality) and 3 (stage-dependent thermal mortality) using calibration with dataset A (Table 4). Circles (**○**) depict dataset A. Filled points (•, ▴) are data not used in calibrations. Triangle (▲) depicts an outlier point.

The yearly thermal and density-dependent mortalities vary by reach and provide information on the spatial/temporal importance of processes that determine juvenile fish survivals. The following results are based on model parameters for Case 5 (Table 4). Figure 7 depicts the partitioning of thermal and density-dependent processes on a reach-specific basis and illustrates that both thermal and non-thermal processes controlled survival but over different intervals of years. Figure 7a illustrates reach-specific survival mediated by thermal mortality and Fig. 7b illustrates the mean reach temperatures at *ATU*crit, which corresponds to *ATU* near the expected hatch. Lower survivals occurred in 2014-2015 when reach temperatures significantly exceeded the critical temperature. Reaches 1 and 2 were mostly below the critical temperature threshold and Reaches 3 and 4 were mostly above the threshold. Reach 4 was above the critical temperature in all years except 2006 (Fig. 7b). In general, the annual variation in density-dependent survival (Fig. 7) was less than the annual variation in temperature-dependent survival (Fig. 7a). The highest density-dependent survivals occurred in Reaches 3 and 4 while the lowest occurred Reaches 1 and 2 (Fig. 7c). This pattern was directly controlled by the redd density in the reaches (Fig. 7d).

**Figure 7.**
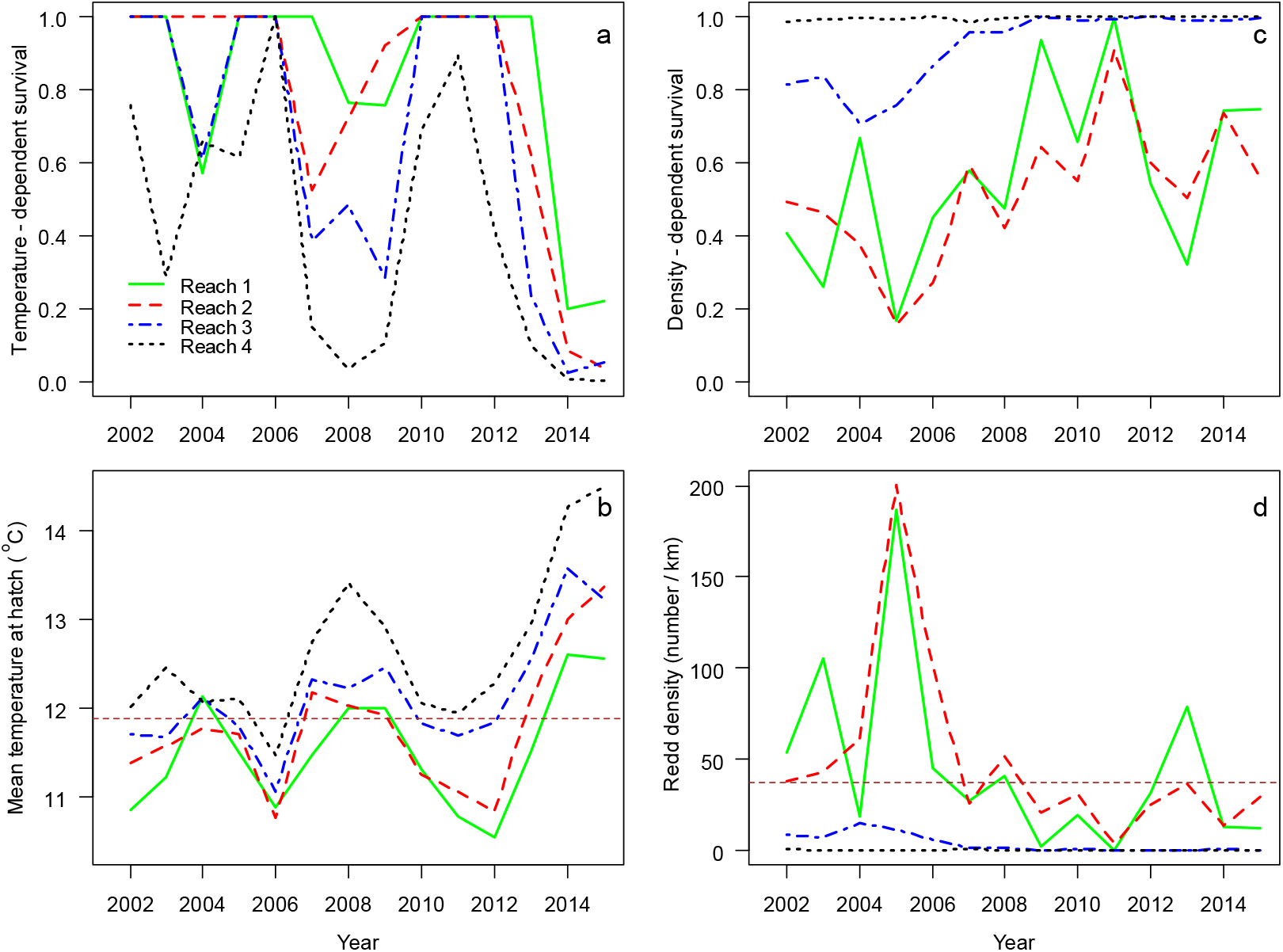
Model Case 5 survival components by year for Reaches 1 − 4: a) mean temperature-dependent survival defined by Eq. (5), b) mean temperature at hatch, c) density-dependent survival defined by Eq. (14), and d) density of redds with carrying capacity. The horizontal dashed lines (---) in 7b and 7d depict critical temperature and carrying capacity respectively.

### Spawning distributions

The motivation of the model was to develop a more mechanistic representation of the effects of temperature control on SRWRC incubation survival. In the BO the water temperature control is designated for May 15 (DOY 135), which can be several weeks prior to spawning and several months prior to the beginning of egg hatching. Thus, pre-season information on spawning is important for modeling reservoir operations. In this section we consider effects of temporal and spatial distributions of redd and assumptions of the critical window on reservoir operations.

### Temporal distribution of incubation stages

Spawn timing involves genetic and environmental factors and is not well predicted. However, the pre-spawning temperature appears to be one factor that affects when salmon spawn. Dusek Jennings and Hendrix (2020) demonstrated that for years 1999-2012 the monthly portions of the total SRWRC spawners were correlated with the April temperature at Keswick Dam. The majority of spawning occurred in either June or July, and using information extracted from their work the fraction spawning in each month, *f*_month_, roughly correlated with April temperature as *f*_*June*_ = 1.6 − 0.11*T*_*april*_ and *f*_*July*_ = −0.83 − 0.12*T*_*april*_. June dominated spawning when April temperature was below 10.5 °C and July dominated when April temperature was above 10.5 °C. Notably, effects of temperature on spawning time have also been observed in other systems. Higher temperatures affected the initiation of upstream migration in the Klamath River (Strange 2010) and temperatures above 13 °C delayed spawning of Pacific Northwest salmon (Richter and Kolmes 2005). For the SRWRC the reach-specific mean spawning day varied by up to six weeks with spawning occurring later upstream by typically a week (Fig 8).

**Figure 8.**
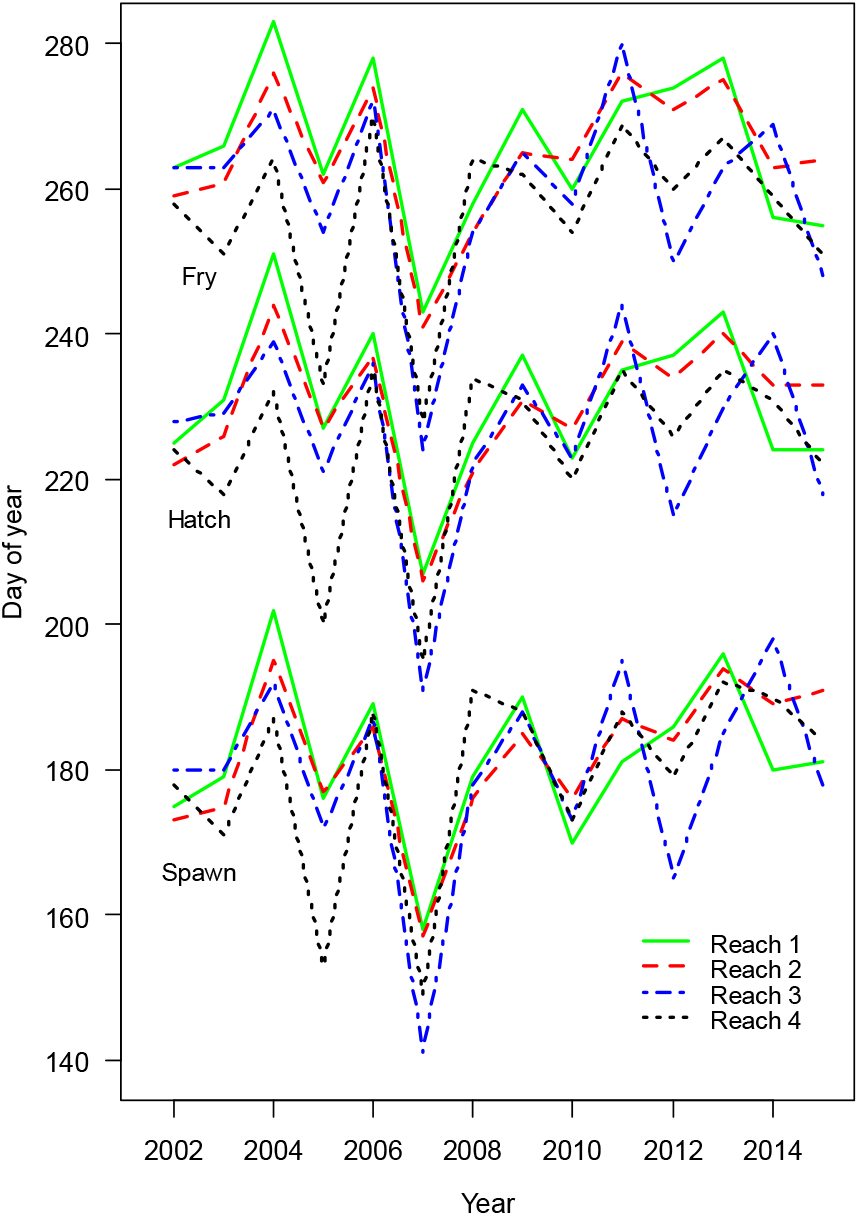
Incubation stage timing by reach and year from Case 5. The hatch date was set using *ATU* = 550 °C d and fry emergence was set using *ATU* = 959 °C d.

If reservoir operations target a critical window, then hatch timing becomes important. This stage is determined by the *ATU* and occurs approximately 40 to 50 days after spawning (Fig. 8). Also, because the upstream reaches are cooler than the downstream reaches, hatching and fry emergence tend to be about a week later in Reach 1 than Reach 3. Thus, river temperature prior to spawning affects spawn timing and temperature after spawning affects the rate of incubation development and hatch timing. Thus, temperatures prior to and during spawning affect fish behavior, which in turn affects the impacts of temperature on fish survival.

### Spatial distribution of redds

In targeting water resources to an incubating population the expected contribution of the reaches to the final population is also important. In particular, in under very dry conditions maintaining a temperature compliance point at Clear Creek may not be compatible with maintaining cool temperatures in Reaches 1 and 2 and so the expected redd distribution may be important for setting the temperature compliance point to a specific reach.

An analysis using Case 5 revealed that the reach-specific distribution of redds and incubation survival were both important in determining the reach-specific contributions to fry passing RBDD (Fig. 9). In general, either Reach 1 or 2 contributed most to fry passage as a result of the centroid of spawning distribution shifting between the two area from one year to the next. Additionally, in the years 2002-2006 the spawning populations were higher and temperatures were lower (Fig. 7) resulting in Reach 3 also contributing to RBDD fry passage in those years. Reach 4, because of its higher temperature and lower spawning numbers, had no significant contribution to the fry population.

**Figure 9.**
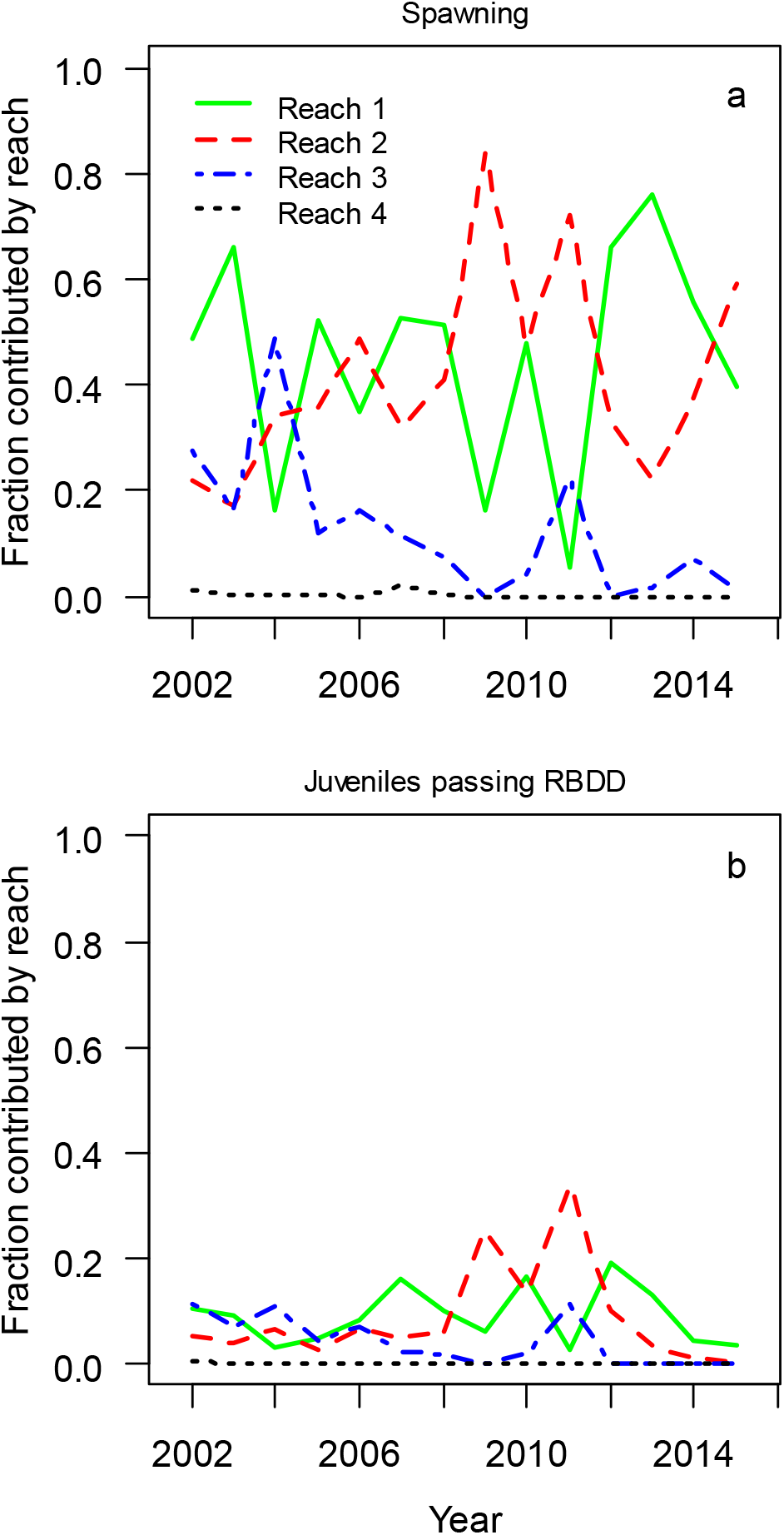
Fractional contributions of reaches for Case 5: a) fractions of redds by reach spawned, and b) fractions of fry by reach passing RBDD.

An analysis of the reach-specific redd locations indicate that the number of spawners in each reach correlated to the total population. In effect, the spawning density in Reaches 1 and 2 increased with the total population size but the relative distribution of spawners between reaches was independent of the population size (Fig. 10). Zero intercept regressions of the reach-specific redd counts against the total numbers of redds gave the reach-specific population fractions *f*_*k*_ of 0.514, 0.334 and 0.188 with r-squares of 0.95, 0.94 and 0.70 for Reach 1, 2 and 3 respectively. These strong relationships indicate that the number of spawners in each reach and year was generally a fixed proportion of the total spawner population. Scaling these fractions by reach length gives an index of reach attractiveness *a*_*k*_ = *f*_*k*_/*L*_*k*_, as the fraction of total redds per unit length of reach. Reaches 1 and 2 have equivalent attractiveness, 0.089, and 0.098 respectively. Reach 3 attractiveness was much less at 0.008. This does not necessarily imply lower attraction to the actual spawning locations in Reach 3. Rather the difference might indicate that effectively, only the upper 2 km of the reach represented desirable habitat. The lower portions might be unsuitable because of temperature or substrate.

**Figure 10.**
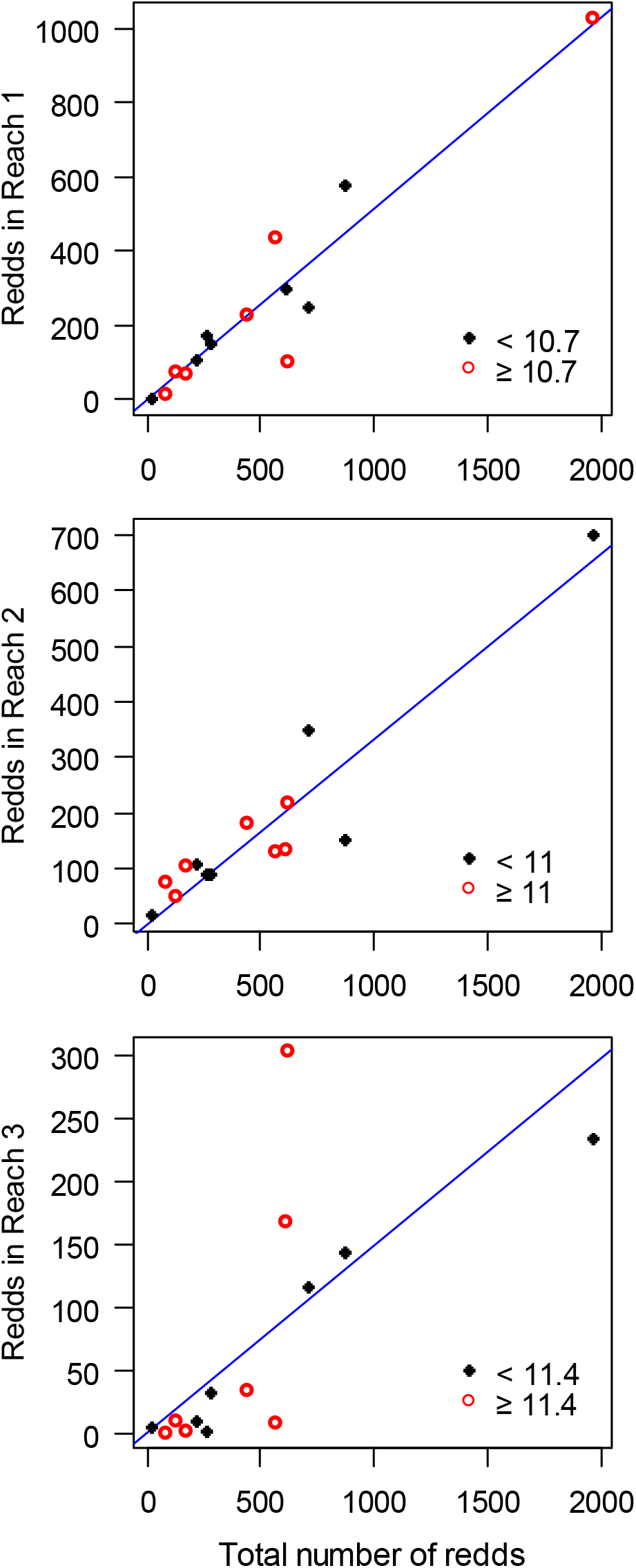
Number of redds in Reaches 1-3 for years 2002-2015. Open points depict temperatures below median for the mean redd date and solid points depict temperatures (°C) above median temperature on the mean redd day. Lines are linear regressions with no intercept.

Figure 10 also characterizes the grand median spawning temperature index, which is the median value of the annual mean spawning temperatures. Importantly, the pattern was not related to Reach 1 and 2 redd counts, suggesting that the spawner distribution was not influenced by an interaction of population size and temperature. However, the figure suggests that few fish occupied Reach 3 in years with small populations and warmer temperatures.

The pairwise comparison of the fractions of redds in the reaches illuminates the possible spawning contagion effect. The fractions of redds in Reaches 1 and 2 laying on the diagonal are inversely related (Fig. 11a) while points falling below the line resided in Reach 3 and are not correlated with the Reach 2 fraction (Fig. 11b). Again, this pattern could be generated by a weakly cohesive spawning group with a center of mass that varied year-to-year between Reach 1 and 2. While the two lowest years were located mostly in Reach 2, the figure again illustrates that the distribution of spawners was not related to population size. Furthermore, neither were the distributions related to spawning temperature. Reach 2 exhibited both low and high fractions of spawners in years with temperatures above and below the grand median spawning temperature. Again, higher spawning temperature might diminish Reach 3 attraction. However, the evidence is weak; a mean fraction of 11% in years with above average temperatures compared to 14% in years with below average temperatures. Additionally, the years with above average temperatures had lower fractions of the population in Reach 3 (Fig. 11b). Thus, the temperature at the mean spawning date appeared to have little effect on the spatial distribution of spawning, except for Reach 3. Evidence for contagion in spawning has also been observed for pink salmon in Alaska (McNeil 1967) and trout in a Minnesota stream (Essington et al. 1998).

**Figure 11.**
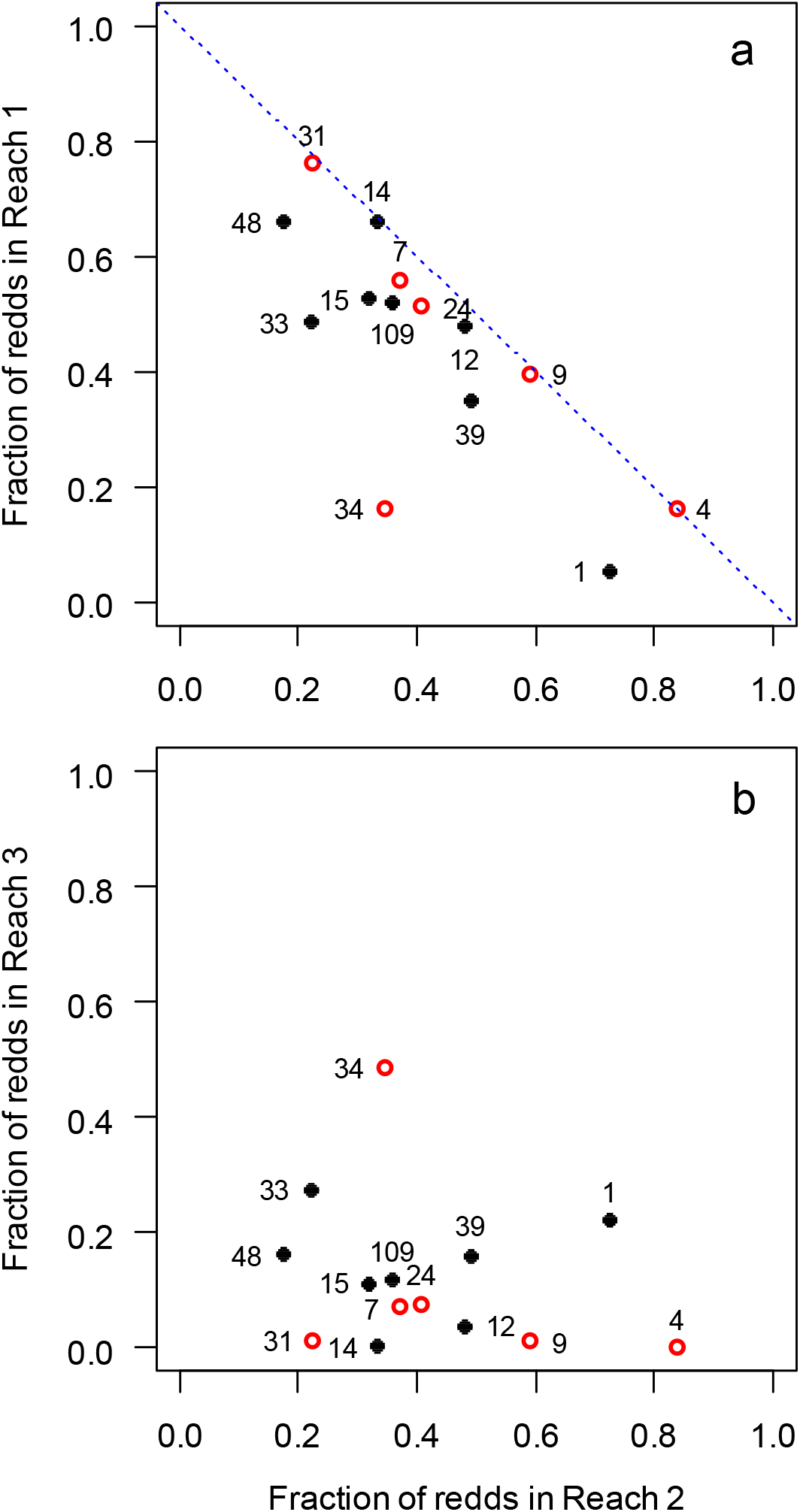
Relationship of population fractions in Reach 2 relative to Reach 1 and Reach 3, years 2002-2015. Numbers depict the ratio of redds in a year to redds in the minimum year 2011, which had 18 redds. Open (closed) points depict years in which the mean spawning temperature was above (below) the across-year mean spawning temperature of 11.76 °C.

Together these observations suggest that the distribution of SRWRC spawning occurred almost entirely in Reaches 1 and 2 with a smaller number spawning in the upper 2 km of Reach 3. Spawning below Clear Creek was negligible. The salient result is that temperature control is most critical in the 9 km stretch of river below Keswick Dam, i.e. to Highway 44 (Table 3). In dry conditions and small spawning populations, the most important spawning habitat is expected to lay within the 9 km length of Reaches 1 and 2.

### Evaluation of reservoir operations

In planning reservoir operations, Reclamation determines the operating tier through several criteria: the cold water pool and total reservoir volume expected in Shasta Reservoir on May 1, flow required to avoid dewatering the redds, flow needed to meet contracted water deliveries for downstream users and the probability of refilling the reservoir the following year. This information guided the development of a cold-water pool management approach based on pre-spawning water availability, or operating Tiers. These pre-season assessments are used to identify which Tier the reservoir is to be operated with to meet the temperature compliance at a temperature gage close to the Clear Creek confluence on the Sacramento River (Fig. 1). The four Tiers, which correspond to decreasing volume in the cold-water pool, are defined:

- Tier 1 –Target 53.5°F (11.9 °C) or lower starting May 15 through October 31
- Tier 2 –Target 53.5 °F during critical egg incubation period and do not exceed 56°F during the management period ending October 31
- Tier 3 – Dry water year targets 53.5-56 °F during critical egg incubation period and do not exceed 56°F during the management period ending October 31
- Tier 4 – Extremely dry water year targets 56 °F (13.3 °C) or higher

Here we evaluate each Tier operation under assumptions that thermal mortality is stage-independent (Case 1) or stage-dependent (Case 3). The simulations were based on the redd distribution in 2013, which represents a year with a relatively smooth bell-shaped distribution of SRWCS. Note that in the comparisons (Table 6) total survivals *S* are higher for Case 1 than for Case 3, while thermal survivals *V* tend to be higher for Case 3. Additionally, Case 3 fits the survival data significantly better than Case 1 (Table 4). Therefore, the comparisons of Tiers in Fig. 12 and Fig. 13 show thermal survivals only. Table 6 also shows the volumes of different temperature waters under the constraint that all Tiers maintained an 8 kcfs reservoir flow, which corresponds roughly with typical Shasta Reservoir spring and summer flows (SRTTG 2018). In dryer years the flow is expected to be less but this variation is not addressed in the example.

**Table 6.** Table illustrates survivals and volumes of water of specific temperatures needed to achieve the Tier compliance temperatures 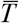. Survivals from spawning to RBDD fry passage for thermal processes only (*V*) and total survival (*S*) for the fish model run with stage-independent and stage-dependent thermal mortality assumptions, Case 1 and Case 3 receptively (Table 4). Storage volume calculated as (temperature step duration) * (8 kcfs reservoir flow). Operation period in all Tiers is June 9 – Oct 17.

| Tier | Storage volumes (KAF) |  |  | Stage-independent model survival |  | Stage-dependent model survival |  |
| --- | --- | --- | --- | --- | --- | --- | --- |
| | $\bar{T} \leq 53.5^\circ$ | $\bar{T} \approx 54.5^\circ$ | $\bar{T} \geq 56^\circ$ | $V$ | $S$ | $V$ | $S$ |
| 1 | 2063 |  |  | 100 | 24.7 | 100 | 19.2 |
| 2a | 2063 |  |  | 96.8 | 24.9 | 100 | 19.2 |
| 2b | 962 |  | 1111 | 71.3 | 18.3 | 93.0 | 17.9 |
| 3a |  | 2063 |  | 97.3 | 25.0 | 99.0 | 18.8 |
| 3b |  | 952 | 1111 | 72.8 | 18.7 | 92.6 | 17.6 |
| 4a |  |  | 2063 | 25.6 | 6.6 | 16.0 | 2.9 |
| 4b |  | 1031 | 1031 | 28.5 | 7.3 | 32.1 | 5.8 |
| 4c | 516 |  | 1547 | 29.4 | 7.6 | 63.0 | 11.5 |

**Figure 12.**
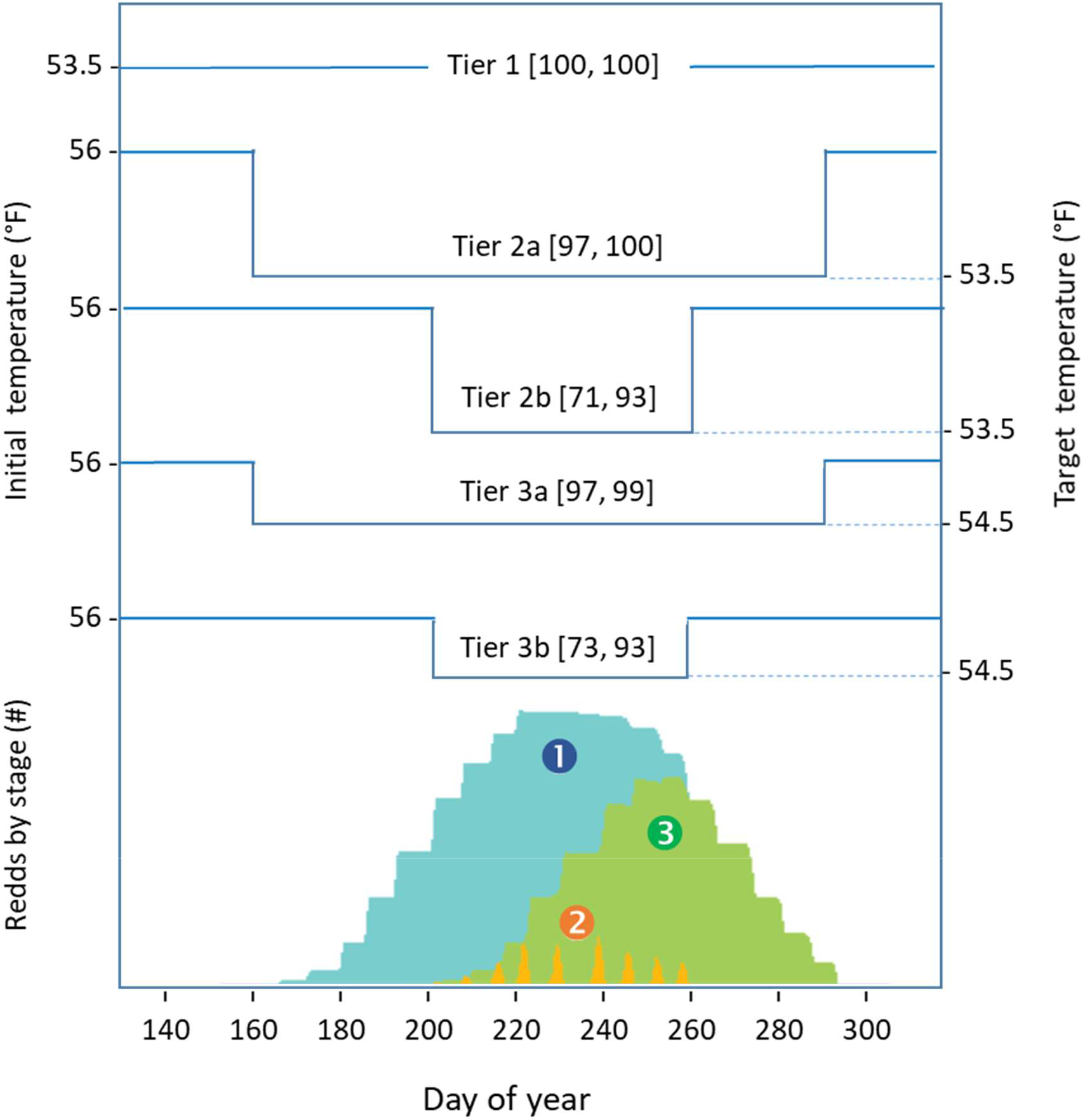
Examples of alternative temperature operations during Above-Normal (Tier 1), Normal (Tier 2), and Below-Normal (Tier 3) water years. Left axis depicts temperature at season beginning, May 15. Right axis depicts temperature targets for critical incubation stage. Bracketed numbers depict percent survivals for stage-independent and stage-dependent thermal mortalities [Case 1, Case 3] (Table 6). Bottom of graph shows number of redds in specific stages by day of year for 2013. Number of redds occupied ➊, number of redds hatching ➋, number of redds with alevin ➌. Transitions in temperature correspond to beginning/end of redd occupancy (Tier 2a, 3a) and beginning/end of hatching (Tier 2b, 3b).

**Figure 13.**
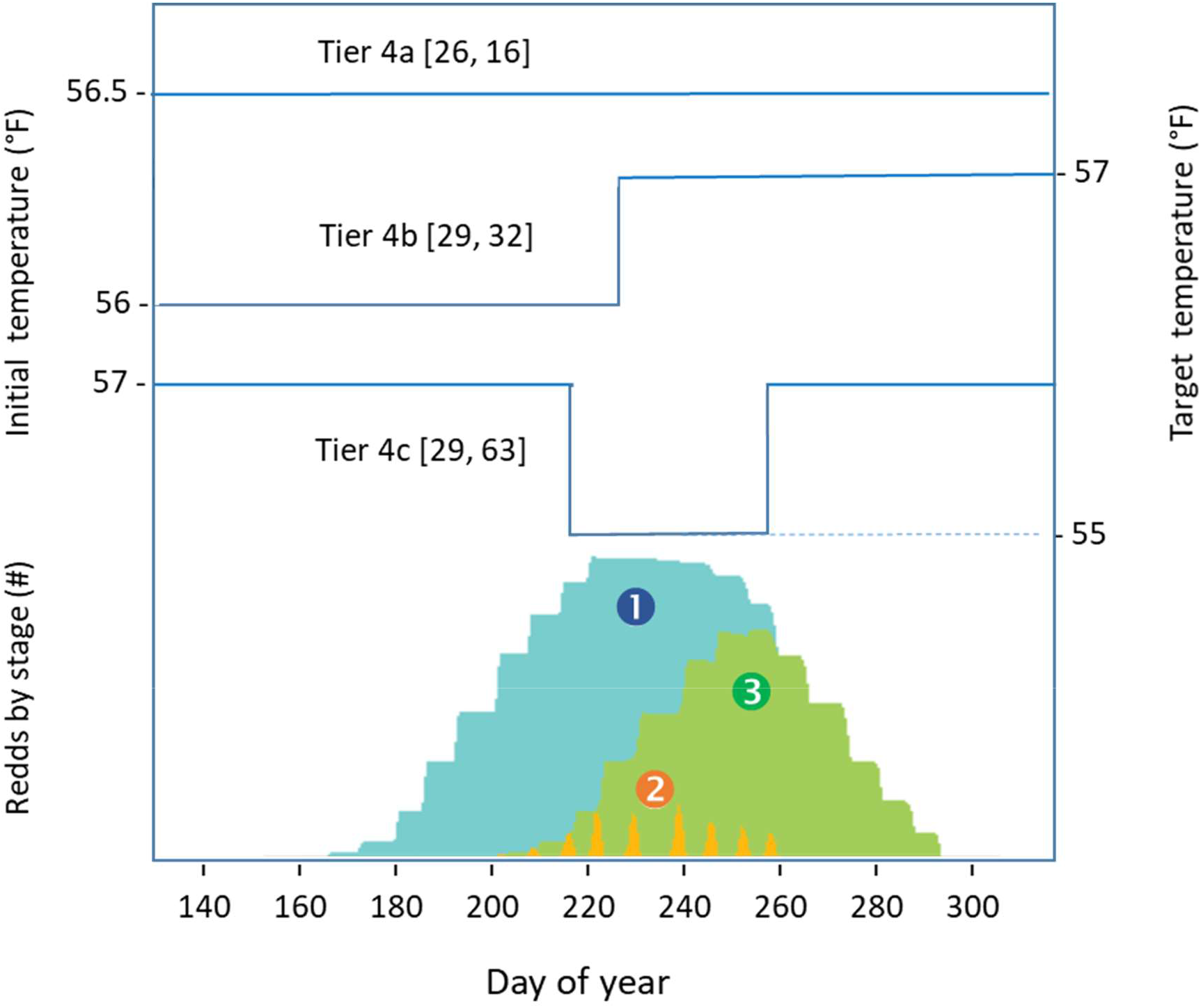
Examples of alternative temperature operations under extremely dry water years (Tier 4). Left axis depicts temperature at season beginning, May 15. Right axis depicts temperature targets for critical incubation stage. Bracketed numbers percent survival from stage-independent and stage-dependent thermal mortalities [Case 1, Case 3] (Table 6). Bottom of graph shows number of redds in specific stages by day of year for 2013. Number of redds occupied ➊, number of redds hatching ➋, number of redds with alevin ➌.

Figure 12 illustrates a range of hypothetical temperature profiles for Tier 1 to Tier 3 conditions with survival calculated for the two critical window assumptions. Tier 1 represents optimum conditions with water temperatures below the critical temperature over the entire incubation. Tier 2a and 3a critical periods correspond to operations under the assumption of stage-independent thermal mortality (Case 1). The cold water delivery covers the entire incubation period, beginning from the creation of the first redd in the season to the emergence of the last fry. Tiers 2b and 3c correspond to operations under the assumption of stage-dependent thermal mortality (Case 3) and extends between the first and last hatches of the spawning season. The first redd-to-last fry period is approximately 60 days longer than the first-to-last hatch period due to summation of hatch time in the first redd and the hatch to alevin emergence time in the last redd.

Figure 13 illustrates Tier 4 corresponding to extreme dry conditions in which the reservoir cold water pool is less than half that available in normal water years. Tier 4a maintains a fixed temperature between June 9 and October 17. Tier 4b temperature starts out low and increases one degree about the middle of the hatch and Tier 4c temperature starts out high and drops two degrees for three weeks around the middle of the hatch. The stage-independent model (Case 1) predicts similar survivals for the three alternatives while the stage-dependent model (Case 3) predicts a factor of four increase in survival between Tier 4a and 4c (Table 6).

## Discussion

In the introduction we noted two timely questions for managing regulated rivers: How much water is needed and When is it needed? (Sabo et al. 2017). In this paper we address these questions for Sacramento River winter-run Chinook salmon through a process-based model of embryo development. The model, designated the Fish Model, was motivated and facilitated by the Endangered Species Act (NMFS 2019b) requirement that Shasta Reservoir cold water storage shall target the incubation stage of the SRWRC salmon. The model predicts incubation survival for temperature control operations under two alternative assumptions of the effect of temperature on incubation mortality: (Case 1) stage-independent mortality occurring over the entire incubation and (Case 3) stage-dependent mortality occurring only over a critical window of embryo development.

With sufficient cold water (Tier 1) both Cases predict the same strategy; control temperature to the lowest practical level across the entire incubation period. However, if the cold water volume is insufficient to maintain the optimal temperature the best strategy depends on the model (Fig. 12, Fig. 13). Under Case 1, the best strategy targets suboptimal temperatures across the entire incubation (Tiers 2a, 3a 4a) but under Case 3, the best strategy targets the optimal temperature to the hatch period (Tier 2b, 3b 4b, 4c). Operating under the wrong assumptions could decrease survival by a factor of two. Additionally, the cold water requirements are significantly different; the duration of Case 1 temperature control is double that of Case 3. Thus, determining which model assumption is most likely to be correct is important for determining reservoir operations.

### Evidence of critical window width

Because of the radically different predictions for the two cases it is important to determine which case is better supported. Our analysis best supports Case 3 in which the critical window occurs near the hatch stage. First, laboratory studies clearly indicate the hatching stage is most sensitive to temperature stress (Rombough 1986). Second, the model estimated *ATU*_crit_ 496 ± 14 °C d corresponds to laboratory studies showing Chinook hatching occurs at 550 ± 122 °C d (Alderdice and Velsen 1978). Third, a short duration stage-dependent thermal window is supported the model fit (Fig. 4, Appendix S2; Fig. S2). The lowest *opt* values occur at δ = 1 to 4 days and the highest at δ ∼ 80 d. Additionally, the lowest ΔAIC scores occur for Case 3, which uses δ = 1 d (Table 4).

Case 3 is also supported through the mechanistic insight it gives to the patterns of model parameters with δ. First, the decreasing *opt* with δ can be view in the context of how the temperature differential Δ*T*_δ_ of Eq. (11) characterizes the mean temperature and thermal stress about *ATU*_crit_. The *opt* exhibits no increase for δ < 4 d and thereafter increases linearly up to δ ≈ 10 d followed by a lessening in the slope. If δ* is an interval centered about *ATU*_crit_, in which embryo response to temperature peaks and the temperatures in the interval are correlated, then the temperature differential Δ*T*_δ_ for any interval δ = δ* ± ε where ε is a small deviation will be similar to Δ*T* resulting in a low *opt*. But a larger δ includes temperature corresponding to the embryo ages not sensitive to temperature. In our analysis this result *opt* increasing for δ > 4. In essence, we reason that the temperature just prior to the hatch realistically characterizes environment and egg condition in which mortality occurs and including additional temperature through larger δ simply adds more noise to the estimation of the thermal differential.

Finally, a short pulse-like critical window of 1 to 4 d comports with oxygen demand quickly rising with just prior to hatch (Fig. 2) and followed by oxygen supply rising at hatch as the embryo switches from cutaneous to branchial respiration (Fig. 3). However, while physiological studies and our model support a pulse-like mortality event for individual eggs, this does not imply that all eggs in a redd experience a coordinated mortality pulse. The heterogeneity within the redd environment (e.g., Hanrahan 2007, Soulsby et al. 2009) could result mortality occurring over multiple days.

### Spawning temperature and location

We note that while δ describes the critical window duration at the scale of the embryo, the model aggregates mortality in all redds and this, in turn, depends on the spatial/temporal distribution of spawners. Thus, at the population-level, the critical window corresponds to the timing of egg hatching, defined by the first and last days of spawning and river temperature. Furthermore, pre-spawning temperature can affect the timing and location of spawning (Dusek Jennings and Hendrix 2020). Consequently, how reservoir operations target the cold water over the incubation season affects both time to hatching and redd locations. For example, higher river temperature at spawning might move the adults up-stream into cooler temperatures but the cooler water could increase the time to hatching. However, higher temperatures might also delay or block spawning (Strange 2010), which from a management perspective might require additional cold water later in the incubation period. These complications were briefly considered in terms of the effect of temperature on the occupancy of the lower regions of the spawning habitat. Overall, the salient point is that pre-spawning temperature likely affects the spawner temporal/special distribution. Thus, targeting river temperature to fish incubation might require iterative modeling over the spawning season as the better information on redds and egg incubation becomes available.

### Density effects

The model formulates density-dependent survival *U* with carrying capacity *D* expressed in redds per river km. This measure is uniquely suited to the SRWRC observation program because redds are indexed by river position, which readily can be aggregated into density. In effect, by incorporating redd location, we obtain a more informative characterization of the mortality processes, as indicated by the better ΔAIC scores for density-based relative to habitat-based population measures (Table 4). Importantly, *B* and *D* estimates covary inversely (Fig. 4) such that higher background survival corresponds with lower carrying capacity. Thus, it is difficult to disentangle these processes using survival data only. However, because of this correlation their combined effect is not influenced by δ nor by uncertainty in *D* (Fig. 5).

### Other uncertainties

While the model fits the data well, r-squares > 0.88 with only temperature and redd density, it disregards factors such as river flow, hyporheic flow ventilation, sedimentation, interstitial biological oxygen consumption, and predation that have demonstrable effects on incubation survival (e.g. Greig et al. 2007, Heywood and Walling 2007, Wildhaber et al. 2014, Cardenas et al. 2016, Sear et al. 2016). We suggest that for SRWRC these other factors are of secondary importance for predicting reservoir operation impacts on fish because the Sacramento River flows are highly regulated during incubation. Thus, sediment accumulation, hyporheic flow and similar processes are expected to have little interannual variability and therefore we consider their contributions as part of the background survival term of the model.

### Real-time reservoir operations

The Shasta Reservoir, as in many highly regulated systems, must operate according to real-time and future commitments that involve multiple temporal scales and changing levels of uncertainty. Reservoirs must maintain storage capacity to capture rain and snowmelt during the winter and spring. Sufficient storage must be reserved for the seasonal temperature needs of critical species while meeting commitments to downstream water users. Thus, effective planning and real-time management necessitates integrating biological, hydraulic and weather models into an integrated system. To this end, the Fish Model, developed in this paper, is implemented as a public web-accessible modeling system as part of the SacPAS website (Fig. 14) (www.cbr.washington.edu/sacramento/fishmodel/). The model draws historical and in-season temperature and spawning information from the SacPAS database (www.cbr.washington.edu/sacramento/data/) and observed and forecast river temperatures from the public website CVTEMP (oceanview.pfeg.noaa.gov/CVTEMP/). The CVTEMP website models water temperature and flow associated with Shasta Reservoir, Shasta Dam operations, and meteorological conditions (Danner et al. 2012, Daniels et al. 2018). The water supply to CVTEMP is obtained from the UCLA’s California and Nevada Drought Monitoring System (http://www.hydro.ucla.edu/monitor_ca/index.html) using information from the NOAA Applied Climate Information System (ACIS) (https://www.rcc-acis.org/).

**Figure 14.**
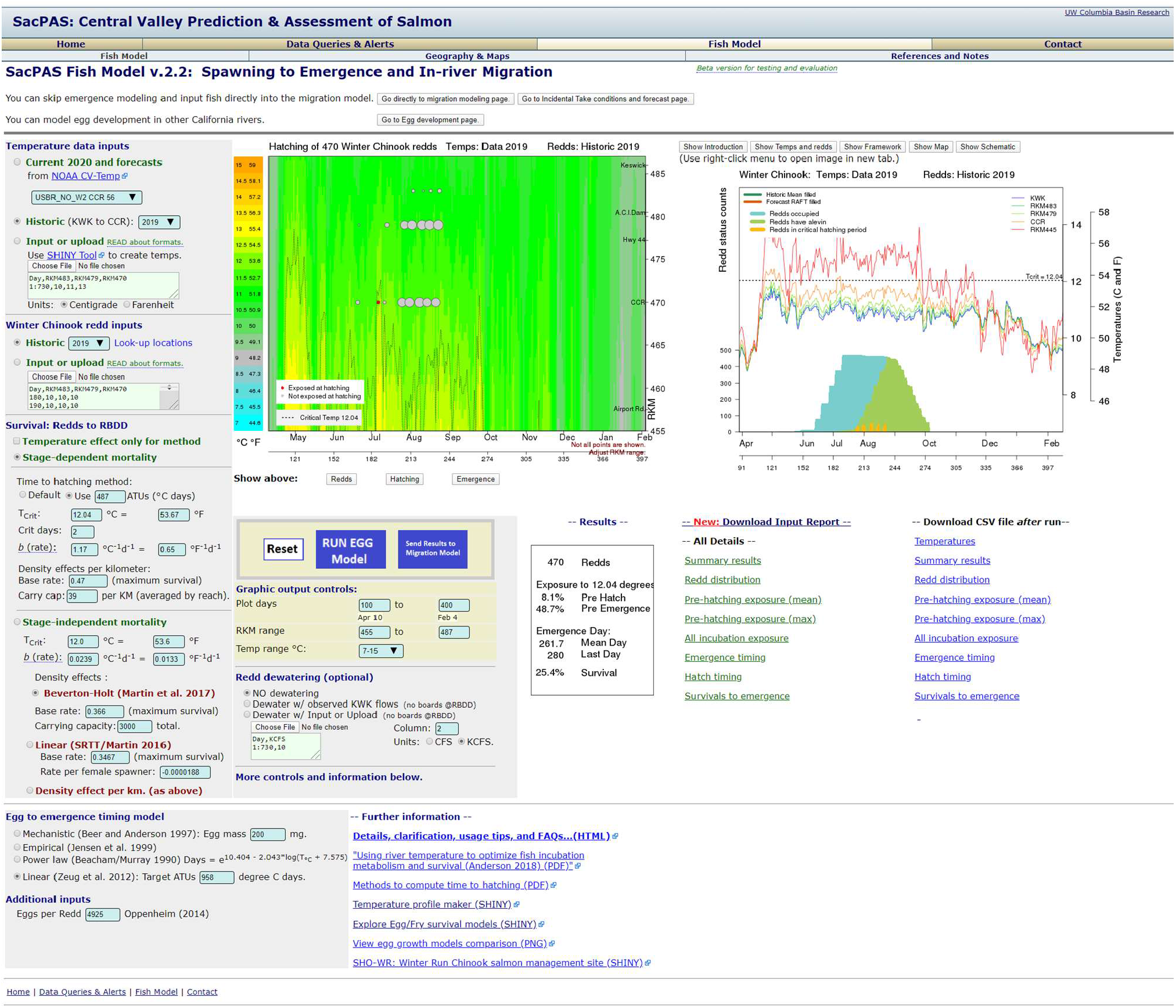
Website of the Egg Model: ➊ data inputs, ➋ stage-dependent thermal mortality configuration, ➌ stage-independent thermal mortality configuration, ➍ egg emergence timing model, ➎ graphical controls and model run, ➏ redd dewatering submodel, ➐ results by date and river km with temperature, ➑ results by stage with redd temperature and stage by date, ➒ results and ➓ all details. Egg Model is available at www.cbr.washington.edu/sacramento/fishmodel/. Historical temperature and spawning data from SacPAS database (www.cbr.washington.edu/sacramento/data/) and forecasts temperature from the NMFS CVTEMP (oceanview.pfeg.noaa.gov/CVTEMP/). (www.cbr.washington.edu/sacramento/data/) and forecasts temperature from the NMFS CVTEMP (oceanview.pfeg.noaa.gov/CVTEMP/).

### Summary

In summary, this paper introduces a mathematical theory for stage-dependent incubation mortality of salmon and demonstrates its importance for efficiently using of cold water resources might be achieved by targeting reservoir operations to the thermally sensitive window of fish incubation. The model was incorporated into and implemented through a web-accessible system developed for the management of Shasta Reservoir for SRWRC salmon. While the model is unique to the Sacramento River, it demonstrates a trend in the collaboration of private, state and federal organizations to develop publicly-accessible web-based models and data for reservoir/river management of threatened and endangered salmon and other species of the California Central Valley (Johnson et al. 2017).

## Acknowledgement

This work was supported by the Bureau of Reclamation contract R17AC00158, and the San Luis and Delta-Mendota Water Authority and Westlands Water District.

## Appendix S1 details of mathematical models

### Relationship of the critical window *δ* to ATU_Y_

To calculate the ATU at *Y* and at the age *y*_crit_ corresponding to the critical window, *ATU*_crit_ the ATU for any age in development is first defined 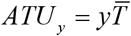 where 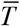 is the mean temperature since fertilization. Next, the mean age in the critical window defined

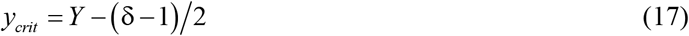

corresponds with *ATU*_crit_. Then *ATU*_crit_ is expressed in terms the mean temperature and either *y*_crit_ or *ATU*_Y_ and δ as

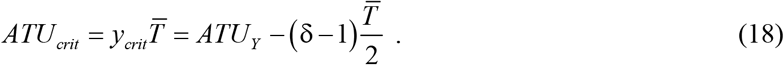

Rearranging Eq. (18) as a linear regression between *ATU*_Y_ and δ gives

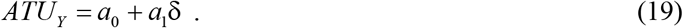

Combining Eq. (19) and (18) then the ATU at the center of the critical window is

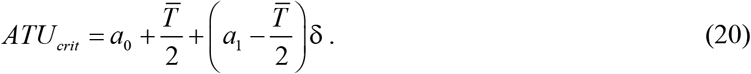

Under the special case of a one-day window, δ = 1 then *ATU*_*crit*_ = *a*_0_ + *a*_1_.

### Relationship of *δ* to the intrinsic rate of thermal mortality b_δ_

To characterize the mortality rate in the critical window first define the sum of daily mortalities within the window for one redd is then

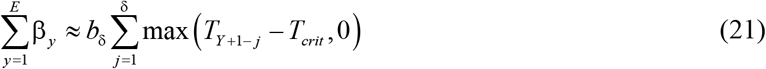

where *T*_*crit*_ is the critical temperature for thermal mortality and *b*_δ_ is a fixed mortality rate coefficient, which depends on δ.

The mean temperature differential in the critical window as

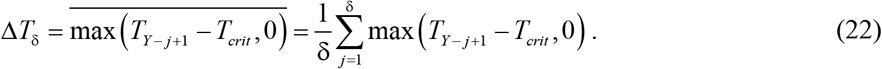

Then the right side of Eq. (21) is approximated *b*_δ_δΔ*T*_δ_. Next the estimation of *b*_δ_ and *T*_crit_ for a given δ involves finding values that minimize the errors in the relation

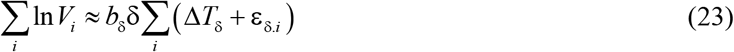

where the index *i* specifies the contribution of the individual redds across all years and the error in the temperature differential is

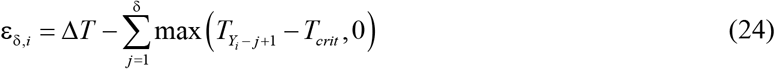

Noting the mortality rate for one day window, i.e., δ = 1, to the rate for any other δ is 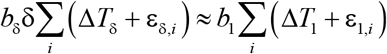 and fitting for each trial δ converges as 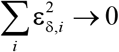 then a regression of *b*_δ_ against 1/δ becomes

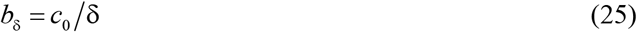

where *c*_0_ = *b*_1_ Δ*T*_1_ / Δ*T*_δ_. However, if Δ*T*_δ_ is independent of δ then *c*_0_ = *b*_1_. This condition was tested in the results by comparing *b*_1_ estimated by fitting the model with δ = 1 against c_0_ determined from a linear regression of Eq. (19) for the model fitted over a range of δ.

### Relationship of *δ* to the model fitting parameter opt

The relationship of the fitting error as a function of δ values is useful for understanding the properties of the critical window. To explore this relationship first assume that Δ*T* is invariant with δ, which implies the mean temperature differential suitably represents the redd thermal exposure across all years. Then from Eq. (23) and (25) the product *b*_δ_δ is constant and the variability in ln*V* depends only on ε_δ,*i*_, which from Eq. (24) is the error in calculating the thermal exposure for a selected δ. Next, assuming no covariance among the errors of the *i* redds and applying the propagation of uncertainty formula, the variance for Eq. (23) is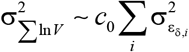.

To characterize the increase of ε_δ *i*_with δ we denote δ* as the best population-level representation of the critical window such that trial values of δ significantly smaller or in particularly significantly larger than δ* produce a critical window with insufficient or irrelevant information. However, the relationship is not simple because it involves both the temperature values within δ and the changing susceptibility of the embryo to temperature relative to *T*_crit_. Both are random processes, so for a simple approximation assume that together temperature and thermal sensitivity with age are characterized by a correlated random walk in which the variance in the thermal mortality rate linearly increases as critical window trials deviate from δ*. We represent this relationship as 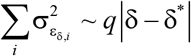 where *q* is the degree of correlation of the terms error for differing critical windows from *j* = 1 to δ in Eq. (24). When the terms in the window are uncorrelated and the process acts as a random walk and *q* = 1. But if terms are completely correlated then *q* = 0. Differing decrees of correlation are then represented by values between 0 and 1 (Renshaw and Henderson 1981). Finally, because *opt*, defined by Eq. (16), monotonically increases with ln*V*, it also monotonically increases with 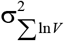 and we link fitting error to the critical window width as

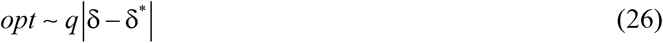

As an aside, management operations of Shasta Reservoir have sought to maintain a stable river temperature during fish incubation. Therefore, the assumption of constant Δ*T* has been facilitated by the temperature control operations.

## Appendix S2

**Appendix S2: Figure S1.**
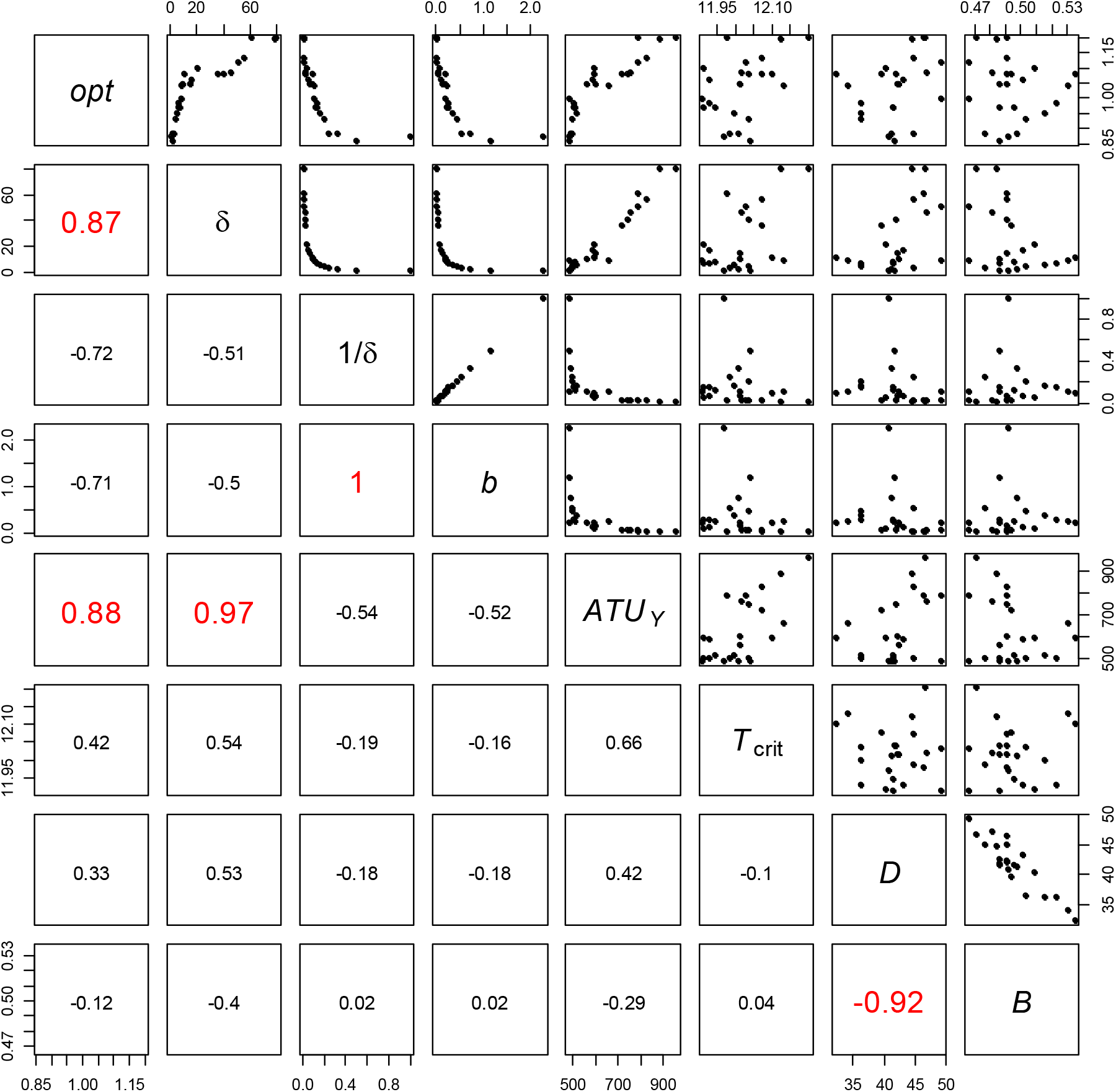
Upper panels show pairwise plots of model parameters for Case 4 using dataset A. Lower panels give correlation coefficients. *T*erms are defined in Table 1.

**Appendix S2: Figure. S2.**
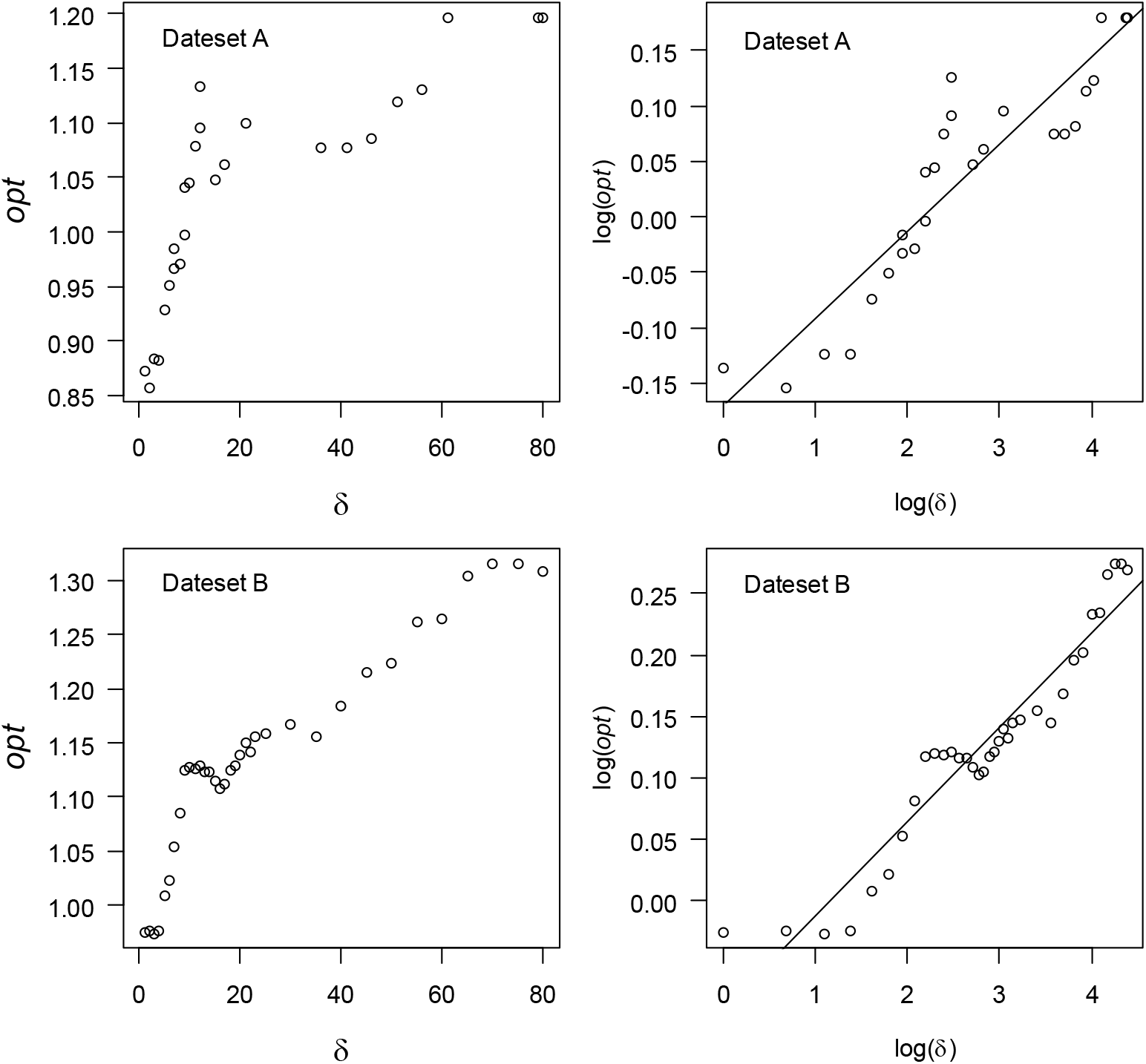
Plots of critical window width δ vs fitting parameter *opt* for datasets A and B.

**Appendix S2: Table S1.** Parameters for Case 4 fit to dataset B for years 1997-1999, 2002-2018. See Table 1 for definitions.

| $\delta$ | $opt$ | r-square | $B$ | $T_{crit}$ | $b_{\delta}$ | $D$ | $ATU_Y$ |
| --- | --- | --- | --- | --- | --- | --- | --- |
| d |  |  | - | °C | 1/°C d | redd/km | °C d |
| 1 | 0.974 | 0.895 | 0.474 | 12.05 | 2.2748 | 43.12 | 482 |
| 2 | 0.976 | 0.893 | 0.478 | 12.04 | 1.0933 | 42.32 | 482 |
| 3 | 0.974 | 0.894 | 0.480 | 12.04 | 0.7434 | 41.81 | 490 |
| 4 | 0.976 | 0.893 | 0.476 | 12.03 | 0.5519 | 41.76 | 500 |
| 5 | 1.008 | 0.893 | 0.484 | 12.02 | 0.4375 | 40.73 | 513 |
| 6 | 1.023 | 0.890 | 0.474 | 12.02 | 0.3570 | 42.26 | 514 |
| 7 | 1.054 | 0.887 | 0.473 | 12.00 | 0.2945 | 44.56 | 514 |
| 8 | 1.084 | 0.889 | 0.474 | 12.01 | 0.2627 | 44.00 | 514 |
| 9 | 1.125 | 0.882 | 0.473 | 12.08 | 0.2489 | 42.86 | 550 |
| 10 | 1.128 | 0.883 | 0.470 | 12.11 | 0.2276 | 42.82 | 575 |
| 11 | 1.126 | 0.881 | 0.474 | 12.11 | 0.2084 | 43.31 | 587 |
| 12 | 1.129 | 0.881 | 0.476 | 12.12 | 0.1905 | 43.16 | 596 |
| 13 | 1.123 | 0.880 | 0.471 | 12.10 | 0.1756 | 43.63 | 602 |
| 14 | 1.123 | 0.880 | 0.476 | 12.12 | 0.1677 | 43.22 | 629 |
| 15 | 1.115 | 0.881 | 0.474 | 12.13 | 0.1568 | 42.93 | 638 |
| 16 | 1.108 | 0.882 | 0.474 | 12.14 | 0.1481 | 43.06 | 650 |
| 17 | 1.111 | 0.883 | 0.469 | 12.16 | 0.1386 | 43.23 | 652 |
| 18 | 1.124 | 0.882 | 0.475 | 12.15 | 0.1303 | 43.20 | 662 |
| 19 | 1.129 | 0.880 | 0.468 | 12.17 | 0.1266 | 44.15 | 690 |
| 20 | 1.138 | 0.884 | 0.473 | 12.15 | 0.1160 | 42.52 | 664 |
| 21 | 1.150 | 0.884 | 0.478 | 12.10 | 0.1069 | 42.43 | 651 |
| 22 | 1.141 | 0.885 | 0.479 | 12.10 | 0.1007 | 42.59 | 651 |
| 23 | 1.156 | 0.883 | 0.477 | 12.08 | 0.0953 | 42.67 | 651 |
| 25 | 1.159 | 0.883 | 0.478 | 12.14 | 0.0907 | 43.01 | 689 |
| 30 | 1.167 | 0.880 | 0.475 | 12.10 | 0.0715 | 44.98 | 708 |
| 35 | 1.156 | 0.883 | 0.477 | 12.05 | 0.0581 | 43.44 | 709 |
| 40 | 1.184 | 0.881 | 0.480 | 12.08 | 0.0516 | 43.33 | 738 |
| 45 | 1.216 | 0.879 | 0.490 | 12.10 | 0.0467 | 42.01 | 785 |
| 50 | 1.223 | 0.877 | 0.483 | 12.14 | 0.0429 | 41.73 | 829 |
| 55 | 1.262 | 0.868 | 0.485 | 12.14 | 0.0385 | 43.67 | 860 |
| 60 | 1.265 | 0.871 | 0.477 | 12.15 | 0.0349 | 43.35 | 860 |
| 65 | 1.303 | 0.861 | 0.475 | 12.21 | 0.0327 | 44.41 | 929 |
| 70 | 1.315 | 0.863 | 0.490 | 12.16 | 0.0298 | 42.48 | 915 |
| 75 | 1.315 | 0.865 | 0.491 | 12.22 | 0.0290 | 41.31 | 943 |
| 80 | 1.308 | 0.864 | 0.476 | 12.20 | 0.0279 | 44.54 | 929 |

**Appendix S2: Table S2.** Parameters of model cases in Table 2 for fits to dataset B and C which is dataset C including the data from 2016. R-sqr. from linear regressions of model survival against data. thermal mortality with critical window extending over entire incubation period. Cases 3 represents stage-dependent thermal mortality.

| Case | Data<sup>1</sup> | $\delta$<br>days | $B$<br>- | $T_{crit}$<br>°C | $b_{\delta}$<br>1/°C d | $D$<br>redd/km | $ATU_Y$<br>°C d | r-sqr |
| --- | --- | --- | --- | --- | --- | --- | --- | --- |
| 3 | C | 79 | 0.439 | 12.13 | 0.0267 | 47.52 | 831 | 0.733 |
| 3 | B | 79 | 0.476 | 12.21 | 0.0291 | 44.33 | 929 | 0.863 |
| 3 | C | 1 | 0.447 | 12.06 | 2.30 | 46.26 | 477 | 0.751 |
| 3 | B | 1 | 0.473 | 12.04 | 2.27 | 43.11 | 482 | 0.895 |

## Notes

### Competing Interest Statement

The authors have declared no competing interest.

